# Endogenous *Syngap1* Alpha Splice Forms Promote Cognitive Function and Seizure Protection

**DOI:** 10.1101/2021.12.05.471306

**Authors:** Murat Kilinc, Vineet Arora, Thomas K. Creson, Camilo Rojas, Aliza A. Le, Julie Lauterborn, Brent Wilkinson, Nicolas Hartel, Nicholas Graham, Adrian Reich, Gemma Gou, Yoichi Araki, Àlex Bayés, Marcelo P. Coba, Gary Lynch, Courtney A. Miller, Gavin Rumbaugh

**Author notes:** Correspondence and Lead Contact: Gavin Rumbaugh, Ph.D. The Scripps Research Institute, 120 Scripps Way, #3B3, Jupiter, FL 33458.

## Abstract

Loss-of-function variants in *SYNAGP1* cause a developmental encephalopathy defined by cognitive impairment, autistic features, and epilepsy. *SYNGAP1* splicing leads to expression of distinct functional protein isoforms. Splicing imparts multiple cellular functions of SynGAP proteins through coding of distinct C-terminal motifs. However, it remains unknown how these different splice sequences function *in vivo* to regulate neuronal function and behavior. Reduced expression of SynGAP-α1/2 C-terminal splice variants in mice caused severe phenotypes, including reduced survival, impaired learning, and reduced seizure latency. In contrast, upregulation of α1/2 expression improved learning and increased seizure latency. Mice expressing α1-specific mutations, which disrupted SynGAP cellular functions without altering protein expression, promoted seizure, disrupted synapse plasticity, and impaired learning. These findings demonstrate that endogenous SynGAP isoforms with α1/2 spliced sequences promote cognitive function and impart seizure protection. Regulation of SynGAP-α expression or function may be a viable therapeutic strategy to broadly improve cognitive function and mitigate seizure.

## Introduction

Pathogenic variation in *SYNGAP1*, the gene encoding SynGAP proteins, is a leading cause of sporadic neurodevelopmental disorders (NDDs) defined by impaired cognitive function, seizure, autistic features, and challenging behaviors [1–8]. *De novo* loss-of-function variants leading to *SYNGAP1* haploinsufficiency cause a genetically-defined developmental encephalopathy (IDC-10 code: F78.A1) that overlaps substantially with diagnoses of generalized epilepsy, global developmental delay, intellectual disability, and autism [4–6, 9, 10]. *SYNGAP1* is completely intolerant of loss-of-function (LOF) variants [11]. Thus, the presence of a clear LOF variant in a patient will lead to the diagnosis of a *SYNGAP1*-mediated developmental encephalopathy. The range of neuropsychiatric disorders causally linked to *SYNGAP1* pathogenicity, combined with the complete penetrance of LOF variants in humans, demonstrate the crucial role that this gene plays in the development and function of neural circuits that promote cognitive abilities, behavioral adaptations, and balanced excitability.

SynGAP proteins have diverse cellular functions [11–13]. The best characterized of these is the regulation of excitatory synapse structure and function located on forebrain glutamatergic projection neurons. In these synapses, SynGAP is predominately localized within the postsynaptic density (PSD), where it exists in protein complexes with synapse-associated-protein (SAP) family proteins [14, 15]. Within these complexes, SynGAP proteins regulate signaling through NMDARs, where they constrain the activity of various small GTPases through non-canonical activity of a RasGAP domain [12, 13]. This regulation of GTPase activity is required for excitatory synapse plasticity [16, 17]. Reduced expression of SynGAP in both human and rodent neurons causes enhanced excitatory synapse function during early brain development and is a process thought to impair cognitive functioning [11, 18, 19]. SynGAP also regulates dendritic arborization. Reduced SynGAP protein expression impairs the development of dendritic arborization in neurons derived from both rodent and human tissues [11, 20, 21], which disrupts the function and excitability of neural networks from both species. While reduced SynGAP expression enhances postsynaptic function regardless of glutamatergic projection neuron subtype, this same perturbation has an unpredictable impact on dendritic arborization, with some neurons undergoing precocious dendritic morphogenesis [11, 20], while others displaying stunted morphogenesis [21]. This is an example of pleiotropy, where *Syngap1* gene products have unique functions depending on the neuronal subtype, or possibly within distinct subcellular compartments of the same type of neuron.

How SynGAP performs diverse cellular functions remains unclear. One potential mechanism is through alternative splicing. Indeed, the last three exons of *Syngap1* undergo alternative splicing[22–24], which results in four distinct C-termini (α1, α2, β, γ). These SynGAP C-terminal protein isoforms are expressed in both rodents and humans, and they are spatially and temporally regulated across mammalian brain development [22, 23]. Moreover, protein motifs present within these differentially expressed C-termini impart SynGAP with distinct cellular functions, with α-derived motifs shown to regulate post-synapse structure and function[25, 26], while the β-derived sequences linked to *in vitro* dendritic morphogenesis [22]. *Syngap1* heterozygous mice, which model the genetic impact of *SYNGAP1* haploinsufficiency in humans, express a robust endophenotype characterized by increased horizontal activity, poor learning/memory, and seizure [12, 16, 18, 27, 28]. Currently, it remains unknown to what extent endogenous *in vivo* expression of alternatively spliced isoforms contribute to systems-level endophenotypes expressed in animal models.

## Results

The last three exons of *Syngap1* undergo alternative splicing **(Fig. 1A)**, which results in four distinct C-termini **(Fig. 1B)**. Exon 19 is spliced into two reading frames (e19b/e19a) **(Fig. 1C)**. Because e19b lacks a stop codon, coding sequences from e20 and e21 are also included in mature transcripts. This leads to expression of α1, α2, or γ C-terminal isoforms **(Fig. 1C-D)**. γ isoforms arise from inclusion e20, while α1 and α2 arise from the absence of e20, but inclusion of e21. e21 itself has two reading frames, with one leading to expression of α1 while the other codes for α2 **(Fig. 1E)**. SynGAP-β arises from splicing of e19 into the “a” reading frame, which contains an internal stop codon **(Fig. 1C)**. To address how expression or function of isoforms contribute to cognitive function, behavior, and seizure latency, we created three distinct mouse lines, each with targeted modifications within the final three exons of the *Syngap1* gene. Each line expressed a unique signature with respect to C-terminal SynGAP protein variant expression or function. For example, in the *Syngap1^td/td^* line, α isoform expression was disrupted while β forms were upregulated **(Fig. 1F-G)**. In contrast, *Syngap1^β*/β*^* mice were opposite with respect to expression of α and β isoforms, with the former upregulated and the later disrupted **(Fig. 1H)**. Finally, the *Syngap1^PBM/PBM^* line, which expressed point mutations that selectively disrupted an essential function of SynGAP-α 1 **(Fig. 1I)**, was useful for determining to what extent phenotypes in the other two lines may have been driven by upregulated or downregulated isoforms.

**Figure 1.**
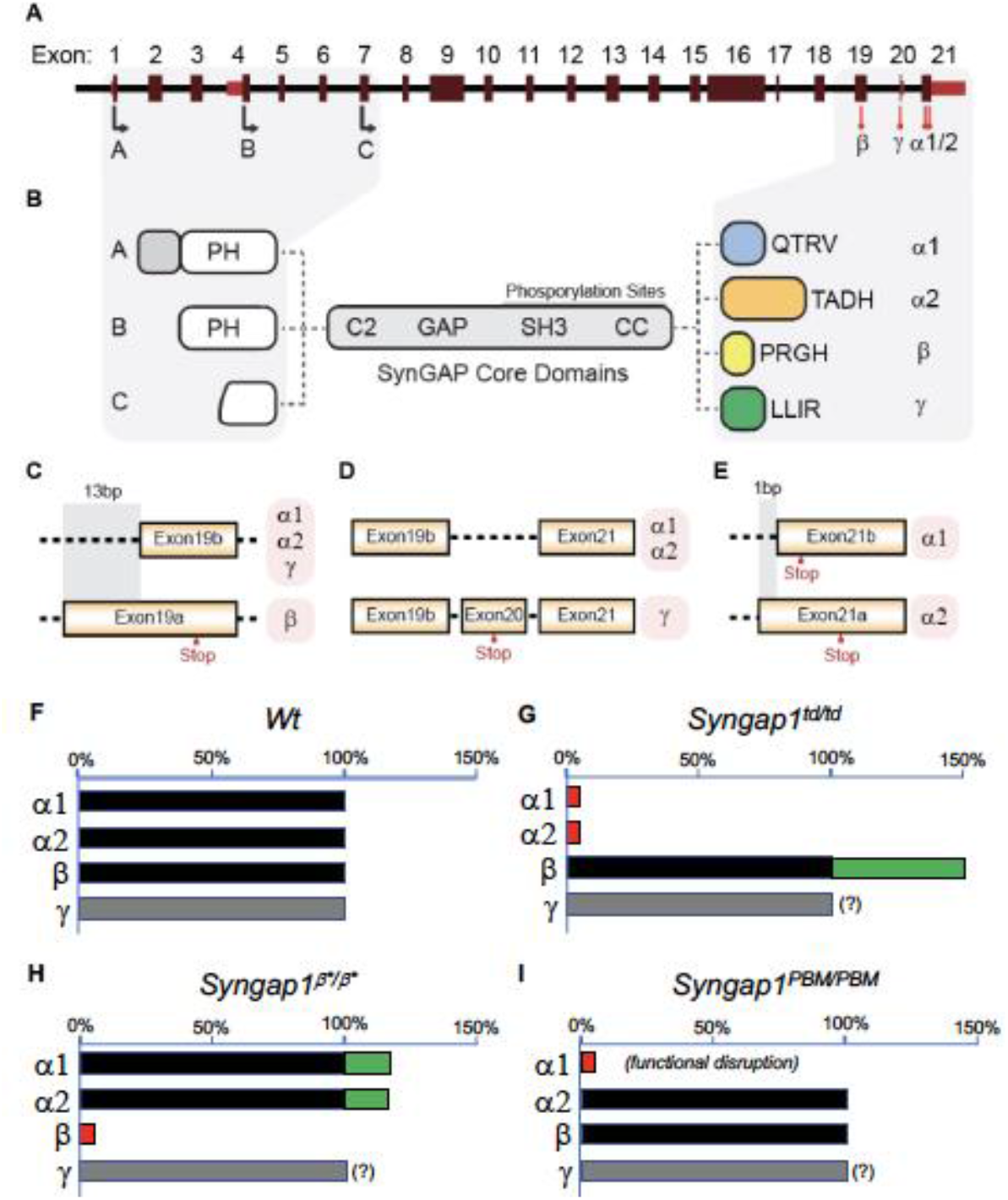
Schematic of *Syngap1 alternative splicing and summary of isoform expression in three new Syngap1 mutant mouse lines*. (A) Map showing alternative use of exons in N- and C-terminal isoforms. N-terminal variants are constituted via use of different start codons in exon1,4 or 7. Exon4 is present only in B-SynGAP. C-terminal isoforms originate from use of different splice acceptors in exon 19 and 21. SynGAP-α1 contains a type-1 PDZ ligand (QTRV). Structure/function relationships of α2, β, γ isoforms remain largely unknown. (B) Schematics of SynGAP isoforms & protein domains. α and β isoforms include full Pleckstrin Homology (PH) domain. In C-SynGAP, this domain is truncated. Core regions common to all isoforms include C2, GAP (GTPase Activating Protein), Src Homology 3 (SH3)-binding, and coiled-coil (CC) domains. Multiple phosphorylation sites are present downstream of the GAP domain. (C-E) Schematics describing C-terminal splicing events producing distinct isoforms. (F-I) Summary of *Wt* and three new *Syngap1* mutant mouse lines each with distinct targeted alleles that disrupt the function or expression of SynGAP C-terminal isoforms. Bars represent expression levels of each C-terminal protein isoform relative to each *Wt* littermate control. Primary data for expression levels can be found in subsequent figures.

### Reduced α1/2 C-Terminal Isoform Expression is Associated with Enhanced Seizure Latency and Cognitive impairment

We previously reported the generation of a *Syngap1* mouse line with an insertion of an IRES-TDtomato (IRES-TD) cassette within the 3’-UTR to facilitate endogenous reporting of active *Syngap1* mRNA translation in cells [29]. The cassette was placed within the last *Syngap1* exon (e21) between the stop codons of α1 and α2 coding sequences **(Fig. 1E; Fig. 2A)**. Our prior study reported neuronal expression of fluorescent protein and normal total SynGAP (t-SynGAP) protein expression as measured by antibodies that recognize all splice forms. Due to our interest in understanding how *in vivo* expression of C-terminal variants impacts brain systems and behavior, we performed an in-depth characterization of behavioral phenotypes and SynGAP isoform expression in IRES-TD mice. Heterozygous (*Syngap1^+/td^*) breeding of IRES-TD animals resulted in offspring of expected mendelian ratios **(Fig. 2B)**. However, while all WT (*Syngap1*^+/+^) mice survived during the 100-day observation period, significant post-weaning death occurred in IRES-TD mice, with approximately two-thirds of homozygous mice (*Syngap1^td/td^*) failing to survive past PND 50 **(Fig. 2B)**. It is well established that complete loss of t-SynGAP protein stemming from homozygous inclusion of null alleles leads to early postnatal death [27, 30]. However, ~50% t-SynGAP expression, like that occurring in heterozygous KO mice (*Figure 2 -supplement 1A*), has no impact on survival [27, 30]. Given the unexpectedly poor survival of *Syngap1^td/td^* animals, we thoroughly examined SynGAP C-terminal isoform protein expression in this line. At PND21, when all three genotypes are abundant **(Fig. 2B)**, t-SynGAP protein in mouse cortex homogenate was reduced in *Syngap1^+/td^* and *Syngap1^td/td^* mice compared to WT controls **(Fig. 2C)**. Reduced t-SynGAP levels appeared to be largely driven by near-complete disruption of α1/2 protein expression from the targeted allele. Reduced α isoform expression coincided with increased protein levels of β-containing C-terminal isoforms. Even with β compensation, *Syngap1^td/td^* mice expressed only ~50% of t-SynGAP at PND21. Whole exome sequencing was carried out in each genotype. Differential gene expression (DGE) analysis revealed only a single mRNA, *Syngap1*, was abnormally expressed (*Supplemental Table 1*). There was a ~25% reduction in mRNA levels in both *Syngap1^+/td^* and *Syngap1^td/td^* mice (*Figure 2 -supplement 1B*). While the IRES-TD cassette destabilized a proportion of *Syngap1* mRNAs, the similarity in mRNA levels from both *Syngap1^+/td^* and *Syngap1^td/td^* samples indicated that other mechanisms must also contribute to reduced protein expression of α1/2 isoforms. Indeed, a recent study identified 3’UTR-dependent regulation of α isoform protein expression [31], suggesting that the IRES-TD cassette is also disrupting translation of these C-terminal variants. We next addressed expression of SynGAP isoforms in adulthood. In this additional experiment, only *Syngap1^+/+^* and *Syngap1^+/td^* mice were used because of limited survival and poor health of homozygous mice in the post-weaning period **(Fig. 2B)**. The general pattern of abnormal SynGAP levels persisted into adulthood, with both α isoforms reduced by ~50% compared to WT levels, while β isoforms were significantly enhanced (*Figure 2 -supplement 1C*). However, the effect on t-SynGAP was less pronounced in older animals and did not rise to significance. This finding highlights the importance of measuring the expression of individual isoforms in addition to total levels of SynGAP protein in samples derived from animal or cellular models.

**Figure 2.**
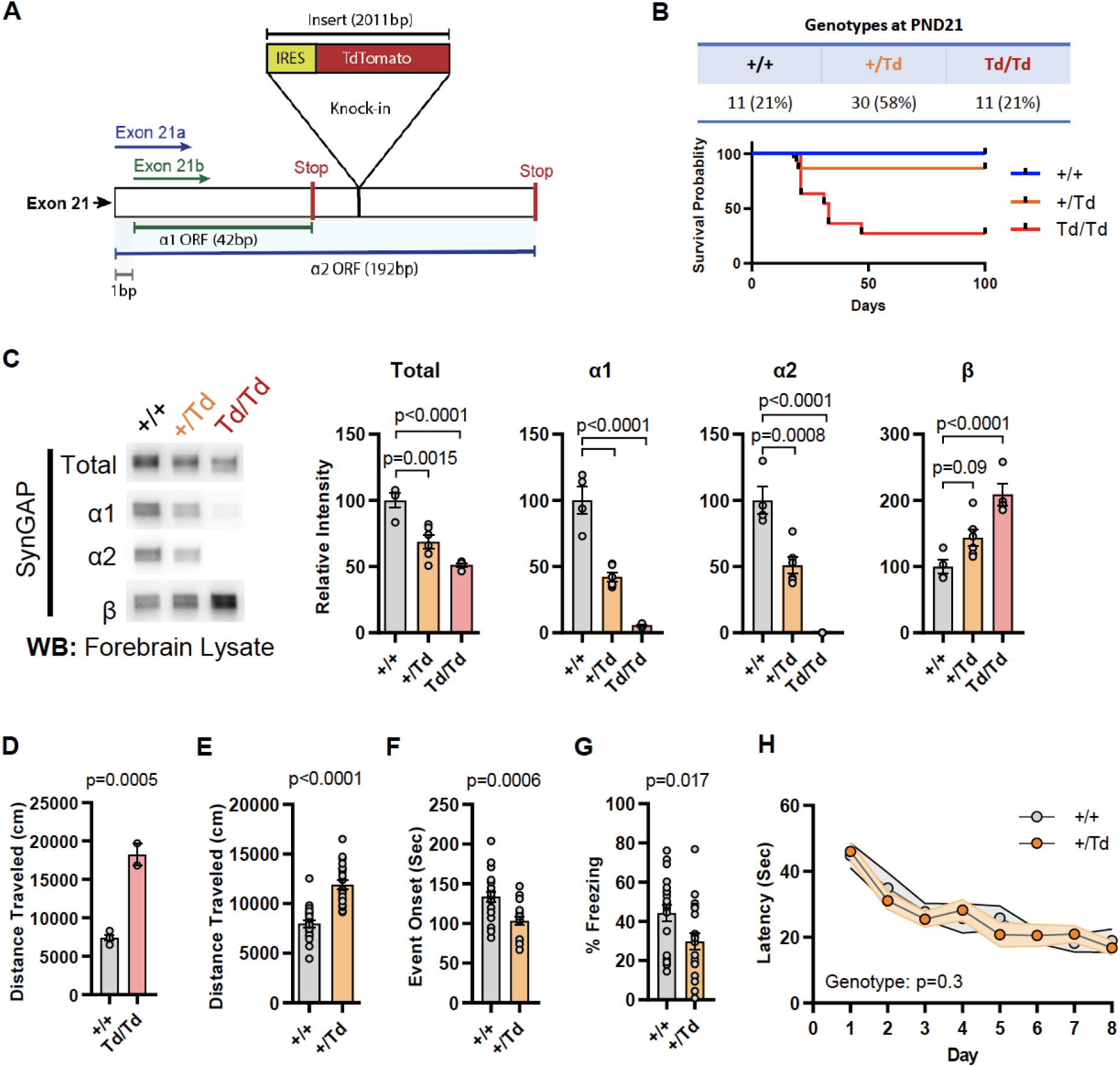
Design and characterization of *Syngap1* IRES-TdTomato knock-in mice. **(A)** IRES-Tdtomato insertion site in relation to α1 and α2 open reading frames. **(B)** Genotype ratios and survival probability following heterozygous breeding. **(C)** Representative western blots showing expression levels of total SynGAP and individual isoforms. Quantification of forebrain expression levels measured by western blot analysis. Relative intensity of bands normalized to total protein signal. Only α1 signal is significantly changed. ANOVA with Tukey’s multiple comparisons test, F(2, 14) = 24.86, n=5, p<0.0001 **(D)** Quantification of total distance traveled in open field test in adult WT or Td/Td mice. Unpaired t-test t(4)=10.42. Note that very few homozygous Td/Td mouse survived through adulthood. **(E)** Quantification of total distance traveled in open field test in adult WT or +/Td mice. Unpaired t-test t(18)=9.007 **(F)** Latency of event onset was measured as the time taken to 1st clonus (seizure onset). Unpaired t-test: t(18)=2.588. **(G)** Percent freezing in remote contextual fear memory paradigm. Unpaired t-test: t(41)=2.49 **(H)** Plots demonstrating latency to find platform across days in Morris Water Maze training. Linear mixed model for repeated measures. n=9-12, +/+ vs +/Td, p=0.3

*Syngap1* heterozygous KO mice, which have 50% reduction of t-SynGAP and 50% reduction of all isoforms (*Figure 2 -supplement 1A*), have normal post-weaning survival rates [27, 30]. However, survival data from *Syngap1^td/td^* mice above, which also expressed a ~50% reduction of t-SynGAP, but loss of α isoform expression **(Fig. 2C; Fig. 1G)**, suggest that expression of these isoforms is required for survival. α isoforms are highly enriched in brain [22], suggesting that reduced survival stems from altered brain function. Therefore, we next sought to understand how reduced α1/2 expression (but in the context of β compensation) impacted behaviors known to be sensitive to reduced t-SynGAP expression in mice. We obtained minimal data from adult *Syngap1^td/td^* mice because they exhibit poor health and survival in the post-weaning period. However, two animals were successfully tested in the open field, and they exhibited very high levels of horizontal activity **(Fig. 2D)**. A more thorough characterization of behavior was carried out in adult *Syngap1^+/td^* mice, which have significantly reduced α isoforms, enhanced β expression, but relatively normal t-SynGAP levels (*Figure 2 -supplement 1A). Syngap1^+/td^* mice exhibited significantly elevated open field activity, seized more quickly in response to flurothyl, and froze less during remote contextual fear memory recall **(Fig. 2E-G)**. These phenotypes are all present in conventional *Syngap1^+/-^* mice [16, 18, 20, 32], which again express ~50% reduction of all isoforms (*Figure 2 -supplement 1A*). In contrast, Morris water maze acquisition, which is also impaired in *Syngap1^+/-^* mice [27, 30], was unchanged in *Syngap1^+/td^* mice **(Fig. 2H)**. Thus, certain behaviors, including horizontal activity, freezing in response to conditioned fear, and behavioral seizure, are sensitive to reduced levels of α isoforms, but not necessarily t-SynGAP levels.

### Enhanced α1/2 C-Terminal Isoform Expression is Associated with Seizure Protection and Improved Cognitive Function

The results in IRES-TD mice suggested that certain core *Syngap1*-sensitive behavioral phenotypes are caused, at least in part, by reduced α1/2 isoform expression. If α isoforms directly contribute to behavioral phenotypes in mice, then increasing their expression may drive phenotypes in the opposite direction. To test this idea, we created a new mouse line designed to upregulate SynGAP-α expression *in vivo*. This line, called *Syngap1^+/β*^* contained a point mutation that prevented use of the e19a spliced reading frame **(Fig. 3A-B)**, the mechanism leading to expression of the SynGAP-β C-terminal variant (**Fig. 1C)**. This design was expected to force all mRNAs to use the e19b reading frame, leading to an increase in α variants (and loss of β expression). This line appeared healthy, bred normally, and resulting offspring were of expected Mendelian ratios (*Figure 3 - supplement 1C*). The CRISPR-engineered point mutation had the predicted impact on SynGAP isoform expression. While there was no change in t-SynGAP expression, there was a copy-number-dependent decrease in β expression, and a modest, but significant, increase in α2 expression in neonatal mice and α1 in young adult mice **(Fig. 3C; Fig. 1H;** *Figure 3 -supplement 1A)*. These animals were then evaluated in behavioral paradigms sensitive to *Syngap1* haploinsufficiency. Homozygous *Syngap1^β*/β*^* mice exhibited significantly less horizontal activity in the open field **(Fig. 3D)**, and also took longer to express behavioral evidence of seizure **(Fig. 3E)**. Further, they expressed no change in freezing levels during remote contextual memory recall **(Fig. 3F).** Unexpectedly, homozygous β* mice exhibited improved learning in the Morris water maze **(Fig. 3G**), with normal memory expression during the probe test (*Figure 3 -supplement 1B*). Thus, a significant increase in α isoform expression (*in the presence of nearly absent β*; **Fig. 1H)** protected against seizure and improved behavioral measures associated with cognitive function, such as learning during spatial navigation.

**Figure 3.**
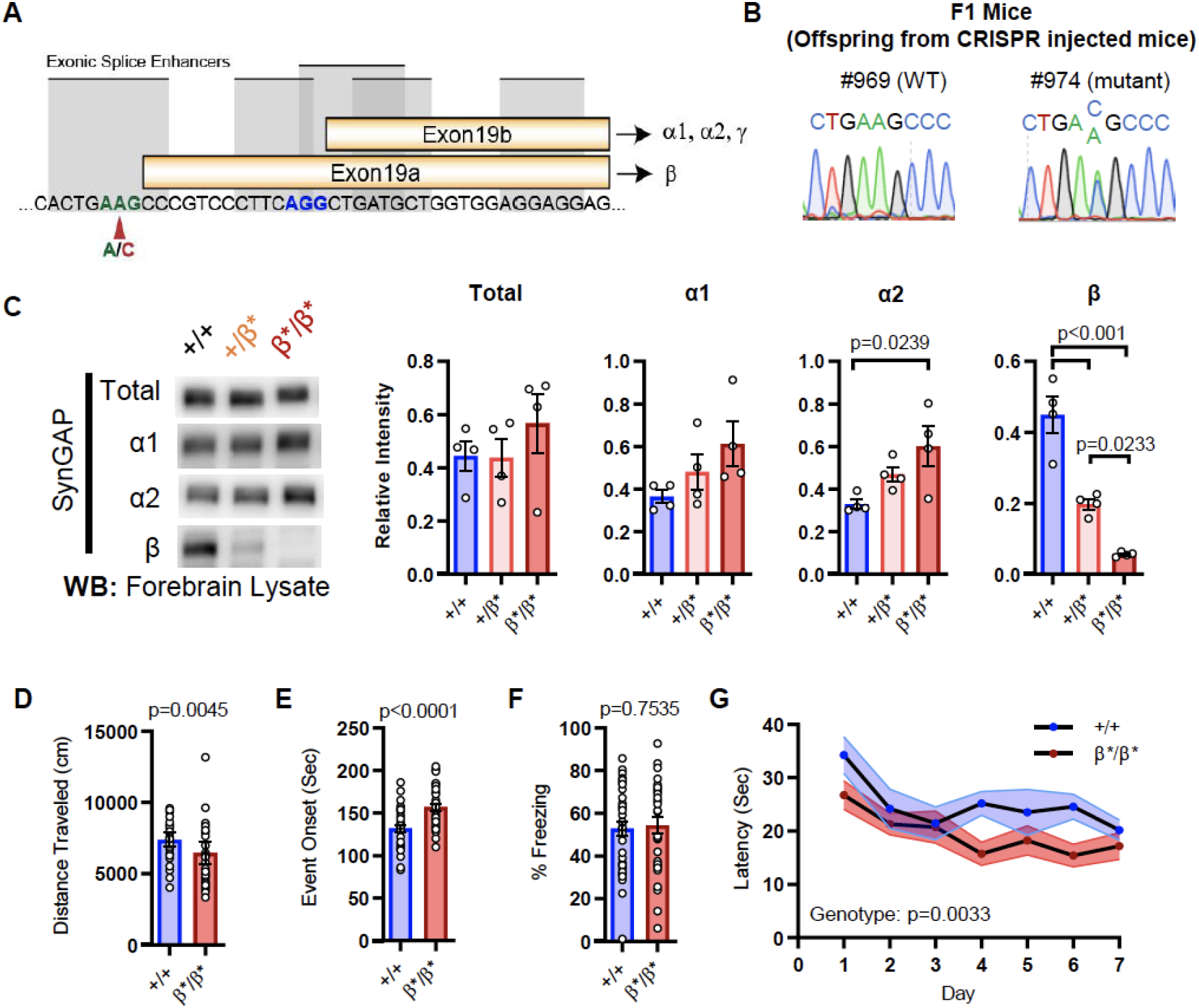
Design and characterization of *Syngap1^β*^* knock-in mice. **(A)** Alternative use of exon19 in distinct splicing events. Exon19 can be spliced into 2 frames shifted by 13 bp. Use of early splice acceptor (green) results in a frameshift and expresses β isoform. Use of the late splice acceptor (blue) allows expression of all other SynGAP C-terminal variants. To specifically disrupt SynGAP-β, a point mutation (A to C) was introduced to the early splice acceptor (indicated with red arrow). **(B)** Sequence trace of *Syngap1^β*/+^* mice obtained via crossing F0 founders to wild-type mice. Mutation site exhibits equal levels of A and C signal in sequence trace indicating heterozygosity. **(C)** Representative western blots showing expression levels of total SynGAP and individual isoforms at PND7. Relative intensity of bands normalized to total protein signal. ANOVA with Tukey’s multiple comparisons test. Total: F(2, 9) = 0.7427, p=0.5029. α1: F(2, 9) = 2.391, p=0.147. α2: F(2, 9) = 5.333, p=0.0297. β: F(2, 9) = 42.53, p<000.1**(D)** Quantification of total distance traveled in OFT. +/+ (n=36), β/β (n=32); Mann-Whitney U=346, p=0.0045. **(E)** Seizure threshold was measured as the time taken to reach three separate events of 1st clonus (event onset) during the procedure. Unpaired t-test t(66)=4.237. **(F)** Percent freezing in remote contextual fear memory paradigm. % Freezing: t(66)=0.3153. **(G)** Plots demonstrating latency to find platform across days in Morris Water Maze training session. Statistical significance was determined by using linear mixed model for repeated measures. Genotype: F(1, 15)=12.22, p=0.0033

Given the observation of seizure protection and improved learning in *Syngap1^β*β*^* mice, we were curious if the impact of the β allele was penetrant in a *Syngap1* heterozygous (*Syngap1^+/-^*) background. This is important given that *Syngap1* heterozygous mice, which model genetic impacts of *SYNGAP1* haploinsufficiency in humans, have seizures and significant cognitive impairments. To test this idea, we crossed *Syngap1^+/β*^* and *Syngap1^-/+^*lines, which yielded offspring with four distinct genotypes: *Syngap1^+/+^, Syngap1^+/β*^, Syngap1^-/+^, Syngap1^-/β*^* **(Fig. 4A)**. We first measured SynGAP protein in each of the four genotypes. In general terms, offspring from this cross expressed changes in SynGAP protein levels that were predicted by the known impact of each allele. For example, the effect of the *Syngap1* null allele (by comparing *Syngap1^+/+^* to *Syngap1^-/+^* samples) was to cause a significant reduction in t-SynGAP, and each of the measured C-terminal isoforms compared to *Syngap1^+/+^* (WT) animals **(Fig. 4B-C,** *Figure 4 - supplement* ). The effect of the *Syngap1* β* allele was to increase both α1 and α2 expression, and decrease β expression, whether the *Syngap1* null allele was present or absent, and these effects were also present at two developmental time points **(Fig. 4B-C,** *Figure 4 - supplement* ). Given these results, we next performed behavioral analyses on all four genotypes. Results on behavioral endophenotypes were consistent with changes in SynGAP protein. For example, the *Syngap1* null allele impaired performance in each of the three behavioral tests performed. Comparing *Syngap1^+/+^* to *Syngap1^-/+^* animals revealed an increase in horizontal distance in the open field, faster time to seizure, and reduced freezing during remote contextual fear recall **(Fig. 4D-F;** two-way ANOVA; null (-) allele, p<0.05). These results replicate many past studies demonstrating the sensitivity of these behaviors to *Syngap1* haploinsufficiency in mice [12, 18, 20, 21, 27, 32, 33]. Interestingly, for both open field and seizure threshold tests, the presence of β* allele significantly improved measures in both WT (*Syngap1^+/+^*) and *Syngap1* heterozygous (*Syngap1^-/+^*) backgrounds **(Fig. 4D-E;** two-way ANOVA; β* allele, p<0.01; interaction of null and β alleles, p>0.5). These findings were consistent with behavioral results from homozygous β* mice in the prior study **(Fig. 3F-G)** and demonstrated that these two behavioral tests are sensitive to the presence of a single β* allele. Also consistent with the prior study in *Syngap1^β*/β*^* mice, the β* allele had no impact on freezing during remote contextual fear recall in either WT or *Syngap1* heterozygous backgrounds **(Fig. 4F)**. Thus, the β* allele partially rescued phenotypes caused by *Syngap1* heterozygosity.

**Figure 4.**
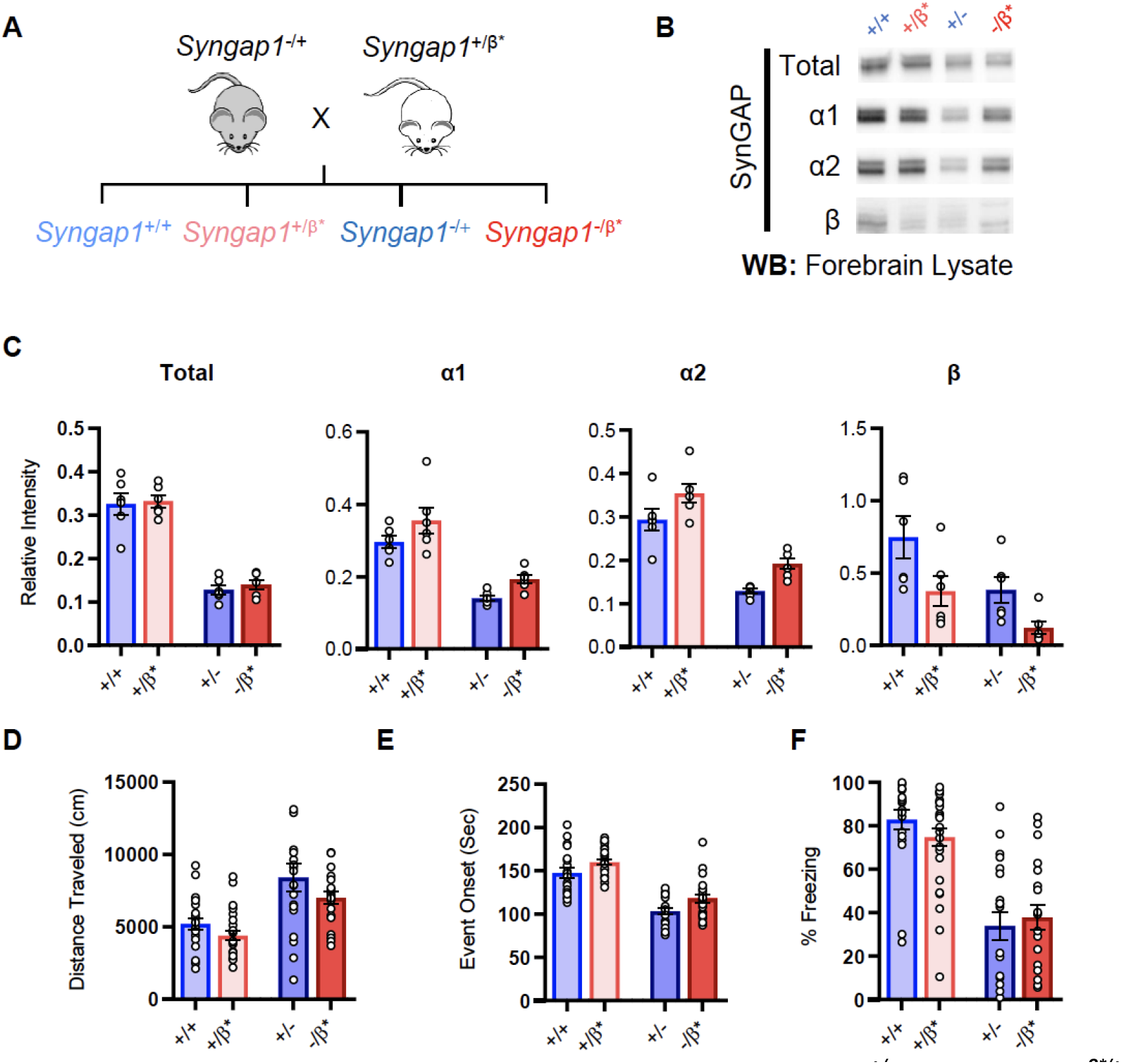
Characterization of offspring derived from *Syngap1^+/-^ and Syngap1^β*/+^* crossbreeding. **(A)** Breeding scheme for offspring genotypes for Syngap1^***+/-***^ and *Syngap1^+/β*^* lines. **(B)** Representative western blots showing expression levels of total SynGAP and individual isoforms at PND7 for all genotypes**. (C)** Quantification of B. Two-way ANOVA with Tukey’s multiple comparison test. **Total:** (-) allele F(1, 20)=146.3, p<0.0001; β* allele F(1, 20)=0.3344, p=0.5696. Allelic Interaction F(1, 20)=0.03191, p=0.8600. **α1:** (-) allele F(1, 20)=56.01, p<0.0001; β* allele F(1, 20)=7.009, p=0.0155; Allelic Interaction F(1, 20)=0.02397, p=0.8785. **α2:** (-) allele F(1, 20)=81.79, p<0.0001; β* allele F(1, 20)=11.92, p=0.0025; Allelic Interaction F(1, 20)=0.0044, p=0.9479. **β:** (-) allele F(1, 20)=9.149, p=0.0067; β* allele F(1, 20)=9.676, p=0.0055; Allelic Interaction F(1, 20)=0.3027, p=0.5883. **(D)** Quantification of total distance traveled in open field test. Two-way ANOVA with Tukey’s multiple comparison test. (-) allele F(1, 86)=28.85, p<0.0001; β* allele F(1, 86)=4.132, p=0.0452; Allelic Interaction F(1, 86)=0.2951, p=0.5884 **(E)** Latency of event onset was measured as the time taken to 1st clonus (seizure onset). Two-way ANOVA with Tukey’s multiple comparison test. (-) allele F(1, 82)=91.71, p<0.0001; β* allele F(1, 82)=8.967, p=0.0036; Allelic Interaction F(1, 82)=0.07333, p=0.7872 **(F)** Percent freezing in remote contextual fear memory paradigm. Two-way ANOVA with Tukey’s multiple comparison test. (-) allele F(1, 86)=69.37, p<0.0001; β* allele F(1, 86)=0.1544, p=0.6953; Allelic Interaction F(1, 86)=1.392, p=0.2414.

### Alpha1 C-Terminal Isoform Function is Required for Cognitive Function and Seizure Protection

The results obtained from *Syngap1* IRES-TD and β* mouse lines indicated that a respective decrease, or increase, in α1/2 isoform expression impaired, or improved, behavioral phenotypes known to be sensitive to *Syngap1* heterozygosity. However, it is also possible that compensatory changes in β expression underlies these phenotypes. This alternative is unlikely, given that α and β expression is anticorrelated in both mouse lines. Thus, for β to drive phenotypes, its expression would need to be both anti-cognitive and pro-seizure, which is inconsistent with isoform expression patterns in *Syngap1^-/+^* mice (*Figure 2 -supplement 1A*), where all protein variants are reduced by half. To directly test the hypothesis that behavioral phenotypes are sensitive to the presence of α isoforms, we attempted to create a third mouse line with point mutations that selectively impacted α isoforms, with minimal effect to SynGAP-β. We took advantage of a known molecular function exclusive to SynGAP-α1. This C-terminal variant is the only isoform that expresses a PDZ-binding motif (PBM). Importantly, cell-based studies have shown that the α1-exclusive PBM imparts unique cellular functions to this isoform [17, 34], such as the ability to become enriched at the post-synaptic density through liquid-liquid phase separation (LLPS). Past studies have shown that mutating the PBM disrupts the ability of SynGAP to regulate synapse structural and functional properties [25, 26], including glutamatergic synapse transmission and dendritic spine size. Before this mouse could be engineered, we had to first identify PBM-disrupting point mutations within the α1 coding sequence that were silent within the open reading frames of the remaining C-terminal isoforms. *In silico* predictions and prior studies [25, 34] suggested that a double point mutation within the α1 PBM could meet these requirements (**Fig. 5A-B)**. To test this prediction, we introduced these point mutations into a cDNA that encoded the PBM and then tested how this impacted PDZ binding. Using an established cell-based assay that reports PDZ binding between the SynGAP PBM and PSD95 [34], we found that these point mutations had a large effect on SynGAP-PDZ binding. When expressed individually in HeLa cells, PSD95-tRFP localized to the cytoplasm, while a SynGAP fragment containing the coiled-coil domain and α1 C-tail (EGFP-CCα1) was enriched in the nucleus **(Fig. 5C-E)**. The co-expression of these two proteins led to SynGAP localization into the cytoplasm. However, this shift in localization did not occur when PBM point mutations were present **(Fig. 5D-E)**, indicating that the selected amino acid substitutions severely impaired binding to the PDZ domains. Moreover, co-immunoprecipitation in heterologous cells indicated that the point mutations in the PBM disrupted the direct association of full-length SynGAP-α1 with PSD95 (*Figure 5 -supplement 1A-B*). Finally, these point mutations also reduced synaptic enrichment of exogenously expressed SynGAP-α1 fragments in cultured forebrain neurons (*Figure 5 -supplement 1C-E*).

**Figure 5.**
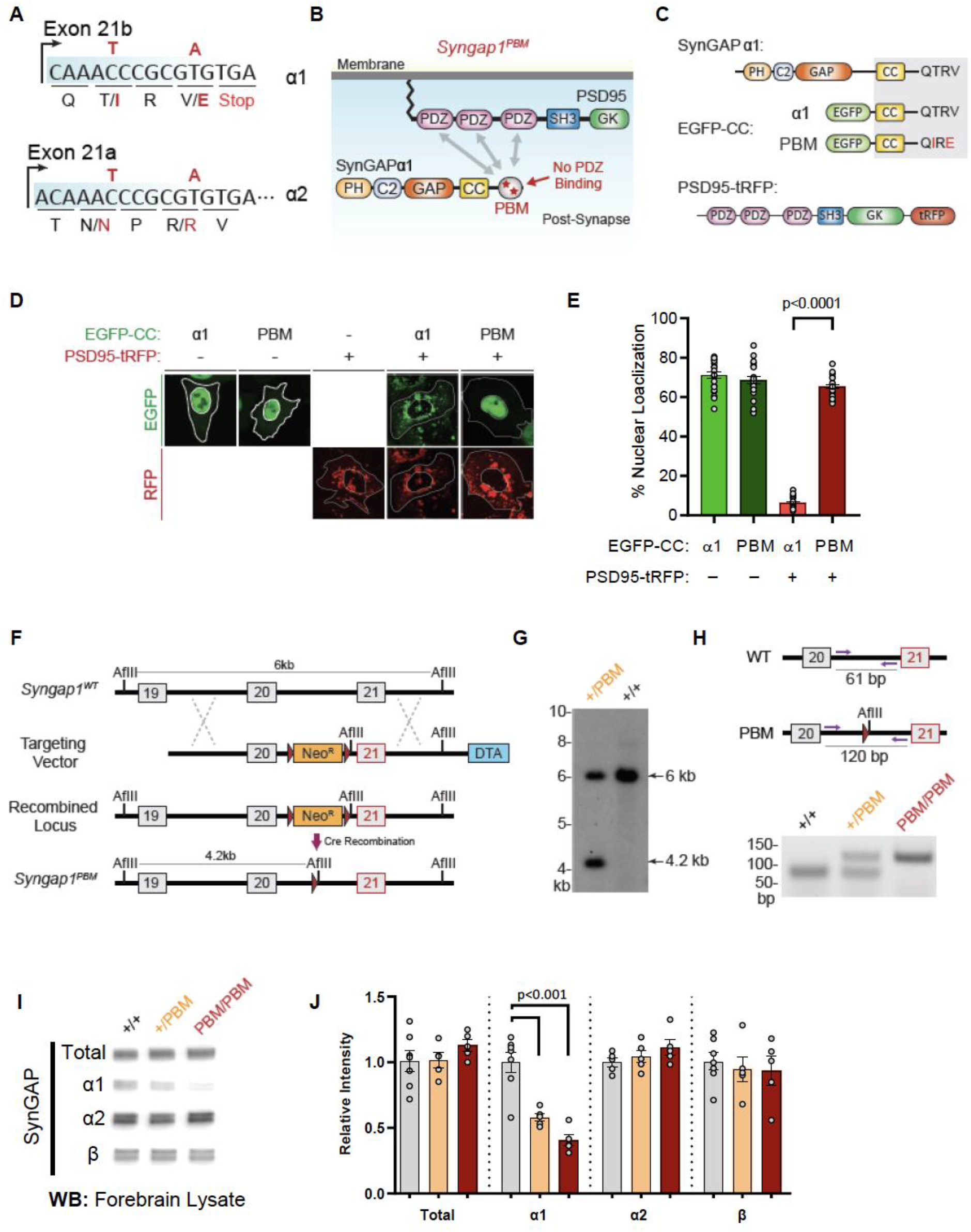
Validation of SynGAP PDZ binding motif (PBM) mutations and construction of the *Syngap1^PBM^* mouse line. **(A)** Schematic diagram for exon map and alternative use of Exon21 in *Syngap1* gene. Exon21b encodes for α1 isoform. Exon 21a encodes for α2 isoform. Point mutations indicated in red alter exon 21b coding sequence without influencing exon21a open reading frame. **(B)** Schematics of SynGAPα1 and PSD95 domain structure and the location of point mutations. **(C)** Illustrations of constructs expressed in HeLa cells to study PDZ-dependent interaction between SynGAP and PSD95. EGFP-CC constructs are homologous to SynGAPα1 C-terminus. **(D)** Co-localization of EGFP-CCα1 and PSD95-tRFP in HeLa Cells. Representative images showing subcellular localizations of WT or PDZ-binding mutant (PBM) EGFP-CCα1 and PSD95-tRFP in HeLa cells when expressed individually or together. **(E)** Quantification of (D). Nuclear localization is calculated as the ratio of EGFP signal colocalized with DAPI vs total EGFP intensity in within an individual cell. ANOVA with Tukey’s multiple comparisons test, F(3, 96) = 531.4. p<0.0001 **(F)** Schematics of the targeting strategy. The targeting vector was spanning Exon20 & 21. The vector included point mutations in Exon21, a neomycin resistance selection cassette flanked by Cre recombination sites and diphtheria toxin selection cassette (DTA). **(G)** Southern blot analysis showing the genomic DNA of the tested heterozygous mice compared to C57BL/6J wild-type DNA. The AflII digested DNAs were blotted on nylon membrane and hybridized with external 5’ probe spanning exon19. **(H)** PCR based genotyping strategy. Primers flanking leftover LoxP site yields 61bp product in WT and 120bp product in mutated allele. **(I)** Representative western blots showing expression levels of total SynGAP and individual isoforms in forebrain lysates. **(J)** Quantification of I. Relative intensity of bands normalized to total protein signal. Only α1 signal is significantly changed. ANOVA with Tukey’s multiple comparisons test, F(2, 14) = 24.86, n=5.

Based on this evidence, we introduced the PBM-disrupting point mutations into the final exon of the mouse *Syngap1* gene through homologous recombination **(Fig. 5A, F-H)**. Both heterozygous and homozygous PBM mutant animals (hereafter *Syngap1^+/PBM^* or *Syngap1^PBM/PBM^*) were viable, appeared healthy, and had no obvious dysmorphic features. We observed Mendelian ratios after interbreeding *Syngap1^+/PBM^* animals (*Figure 5 -supplement 1F)*, demonstrating that disrupting the PBM had no impact on survival. Western blot analysis of forebrain homogenates isolated from *Syngap1^+/PBM^* or *Syngap1^PBM/PBM^* mutant animals demonstrated no difference in t-SynGAP protein levels using antibodies that detect all SynGAP splice variants **(Fig. 5I-J)**. Moreover, using isoform-selective antibodies [35], we observed normal expression of SynGAP-β and SynGAP-α2 isoforms **(Fig. 5I-J).** A reduced signal of ~60% was observed in samples probed with α1-specific antibodies. However, we also observed a similarly reduced signal in heterologous cells expressing a cDNA encoding the mutant PBM (*Figure 5 -supplement 1G-I*), indicating that these antibodies have reduced affinity for the mutated α1 motif. Together, these data strongly suggest that the α1 variant is expressed normally in *Syngap1^PBM/PBM^* animals. This interpretation was supported by RNA-seq data, where normal levels of mRNA containing the α1 reading frame were observed in brain samples (*Figure 5 -supplement 1J*). These data, combined with the observation of no change in total SynGAP protein expression in *Syngap1^PBM/PBM^* samples **(Fig. 5I-J)**, strongly support the conclusion that the PBM-disrupting point mutations do not change the expression levels of the major SynGAP C-terminal splice variants, including those containing the PBM. Thus, this animal model is suitable for understanding the putative biological functions mediated by α1-specific splicing.

Given the disruption to SynGAP-α1 PBM, we sought to understand how disrupting this functional motif impacted previously defined features of SynGAP at excitatory postsynapses. α1 is believed to be anchored within the PSD in part through PBM binding to PDZ domain containing proteins. However, SynGAP molecules multimerize in vivo and it is currently unknown if this results in homo- or hetero-multimerization. Thus, it is unclear how a functional disruption to one isoform generally impacts native SynGAP complexes at synapses. t-SynGAP levels were reduced in PSD fractions prepared from the hippocampus of *Syngap1^PBM/PBM^* mice **(Fig. 6A)**. Importantly, a corresponding increase in t-SynGAP was observed in the triton soluble synaptosomal fraction in these mice, further supporting the observation of reduced t-SynGAP levels in the PSD. We observed similar reductions in t-SynGAP levels within the PSD and ERK1/2 signaling was elevated in neurons cultured from *Syngap1^PBM/PBM^* mice **(Fig. 6B)**. Acute treatment with the NMDAR antagonist APV normalized SynGAP levels in both PSD preparations and normalized ERK1/2 phosphorylation **(Fig. 6B)**. Similar treatments also normalized enrichment of SynGAP in dendritic spines and surface expression of GluA1 in neurons derived from *Syngap1^PBM/PBM^* mice **(Fig. 6 C, D).** These results indicate that endogenous PBM binding regulates an NMDAR-dependent process within excitatory synapses.

**Figure 6.**
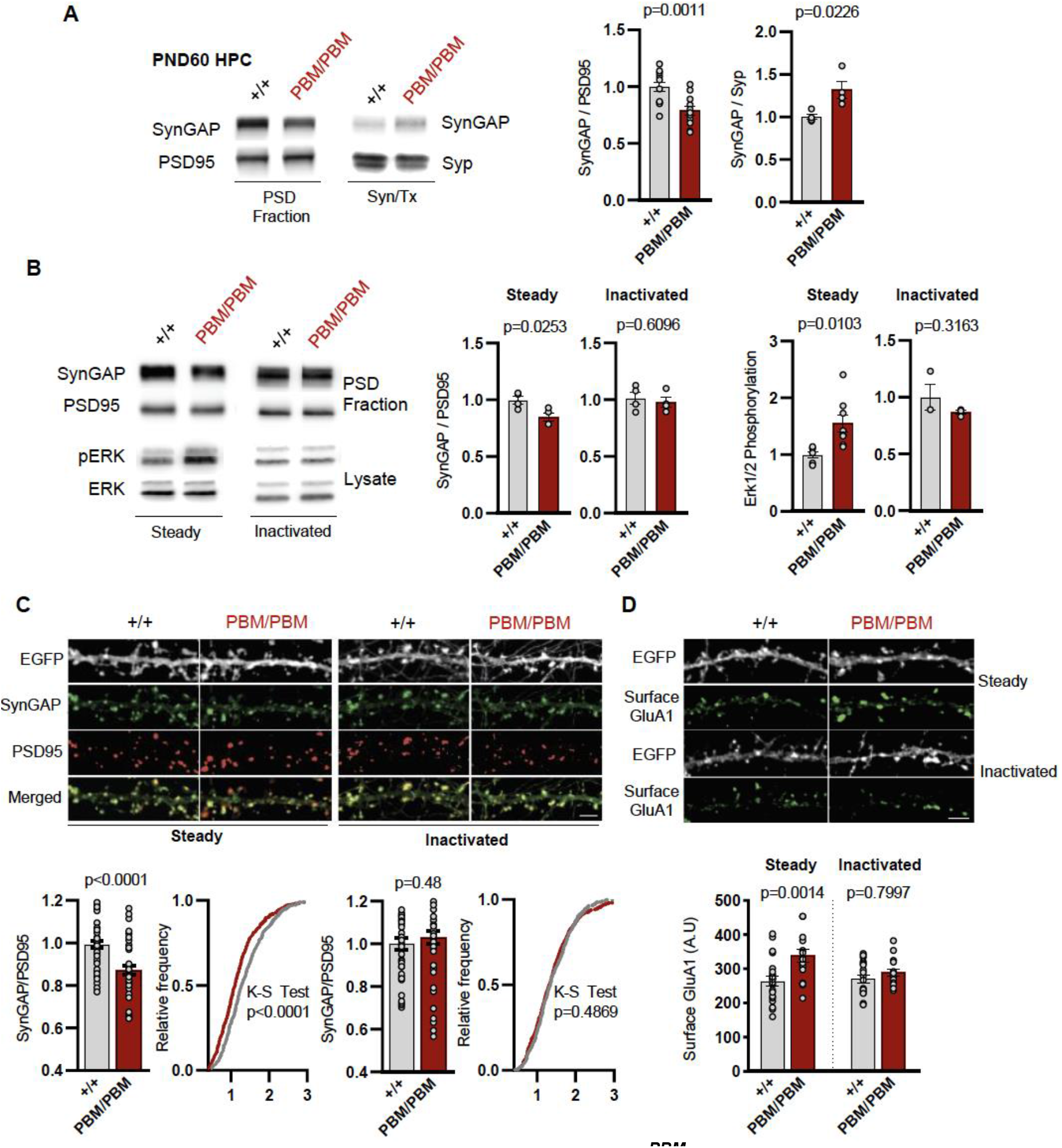
SynGAP synapse localization in *Syngap1^PBM^* mouse line. **(A)**Western blots showing relative distribution of SynGAP in PSD and Syn/Tx fractions from adult hippocampi. Quantification of western blots probing total SynGAP, Synaptophysin and PSD95. For PSD fractions PSD95 and for Syn/Tx fractions Synaptophysin (Syp) were used as loading control. PSD fractions: t(22)=3.733, p=0.0011 n=12 (3 technical replicates for each sample), Syn/TX fractions: t(6)=3.049, p=0.0226, n=4. Each sample represents hippocampi pooled from 2 mice. **(B)** Western blots showing relative enrichment of (i) SynGAP and PSD95 in PSD fractions isolated from DIV18-21 cultures, (ii) phospho and total-ERK1/2 levels in whole cell lysates in steady or inactivated state. Synaptic enrichment of SynGAP in (i) steady-state: Unpaired t-test, t(12)=3.040 p=0.0103. (ii) inactivated state: Unpaired t-test, t(6)=0.5385 p=0.6096. Erk1/2 phosphorylation is calculated as ratio of phospho-Erk1/2 to total-Erk1/2 in homogenates. Erk1/2 phosphorylation in (i) steady-state: Unpaired t-test, t(6)=2.961 p=0.0253. (ii) inactivated state: Unpaired t-test, t(4)=1.144 p=0.3163**(C)** Synaptic enrichment of total SynGAP in WT and PBM mutants in steady or inactivated state. Levels of SynGAP relative to PSD95 signal in dendritic spines. Left, bar graphs demonstrate mean enrichment in an individual dendritic segment. Steady-state: t(90)=4.393 p<0.0001. Inactivated: t(78)=0.6982 p=0.48. Cumulative distribution of SynGAP to PSD95 ratios in individual synapses. Kolmogorov-Smirnov test, Steady-state: p<0.0001, Inactivated: p=0.4869. **(D)** Surface GluA1 expression in primary forebrain cultures in steady or inactivated state. Quantification of mean surface GluA1 levels coincident with PSD95 puncta. Two-way ANOVA with Tukey’s multiple comparisons test. Interaction: F(1,74)=4.112, p=0.0462, Genotype: F(1,74)=11.09, p=0.0014. Treatment: F(1,74)=2.329, p=0.1313. Each n represents an average of 25-30 spines from a dendritic segment belonging to distinct neurons.

Blocking NMDAR activity in *Syngap1^PBM/PBM^* neurons prevented alterations in SynGAP levels at postsynapses **(Fig. 6A-D)**. This suggested that the PBM regulates SynGAP-specific functions in the PSD. However, SynGAP-α1 undergoes LLPS and this mechanism is thought to facilitate the organization of the PSD [34]. Thus, disrupted SynGAP post-synaptic levels could also be attributable to altered structural organization of the PSD. To determine if the PBM contributes to the organization of macromolecular complexes within excitatory synapses, we immunoprecipitated PSD95 from neurons obtained from either WT or *Syngap1^PBM/PBM^* mutant neurons. These neurons were treated with APV to avoid the confounds of elevated NMDAR signaling. These samples were then analyzed by mass spectrometry to determine how disrupting SynGAP-PDZ binding impacted the composition of PSD95 macromolecular complexes. In general, we found only minor differences in the abundance of proteins that comprise PSD95 complexes when comparing samples from each genotype **(Fig. 7A)**. Only 1 out of ~161 proteins (from 133 distinct genes) known to be present within PSD95 complexes [36] met our threshold for significance, although there were modest changes in proteins with structurally homologous PBMs (Type-1 PDZ ligands), such as Iqseq2 and Dlgap3 **(Fig. 7B)**. However, the vast majority of related PBM-containing proteins were not different in mutant neurons, including NMDAR subunits and TARPs **(Fig. 7C)**. Consistent with the mass spectrometry analysis, immunoblot analyses found no changes in TARPs or LRRTM2 in isolated PSDs from *Syngap1^PBM/PBM^* mice **(Fig. 7D-G)**. Although PDZ binding was disrupted, SynGAP protein levels were also unchanged within PSD95 complexes, a result consistent with PSD and synapse localization measurements in APV-treated neurons derived from *Syngap1^PBM/PBM^* mice **(Fig. 6B-C).** These results indicate that SynGAP interacts with PSD95 in a non-PDZ-dependent manner. In support of this interpretation, there is significant overlap between the interactomes of PSD95 [36] and SynGAP [37] macromolecular complexes **(Fig. 7H)**. Thus, within intact postsynapses, SynGAP and PSD95 interact, as part of a macromolecular complex, through binding to common protein intermediaries. Together, these data suggest that SynGAP PBM binding to PDZ domains is not a major factor promoting the organization of PSD95 macromolecular complexes or the PSD. Rather, the PBM appears to regulate SynGAP-specific mechanisms that control signaling through NMDARs.

**Figure 7.**
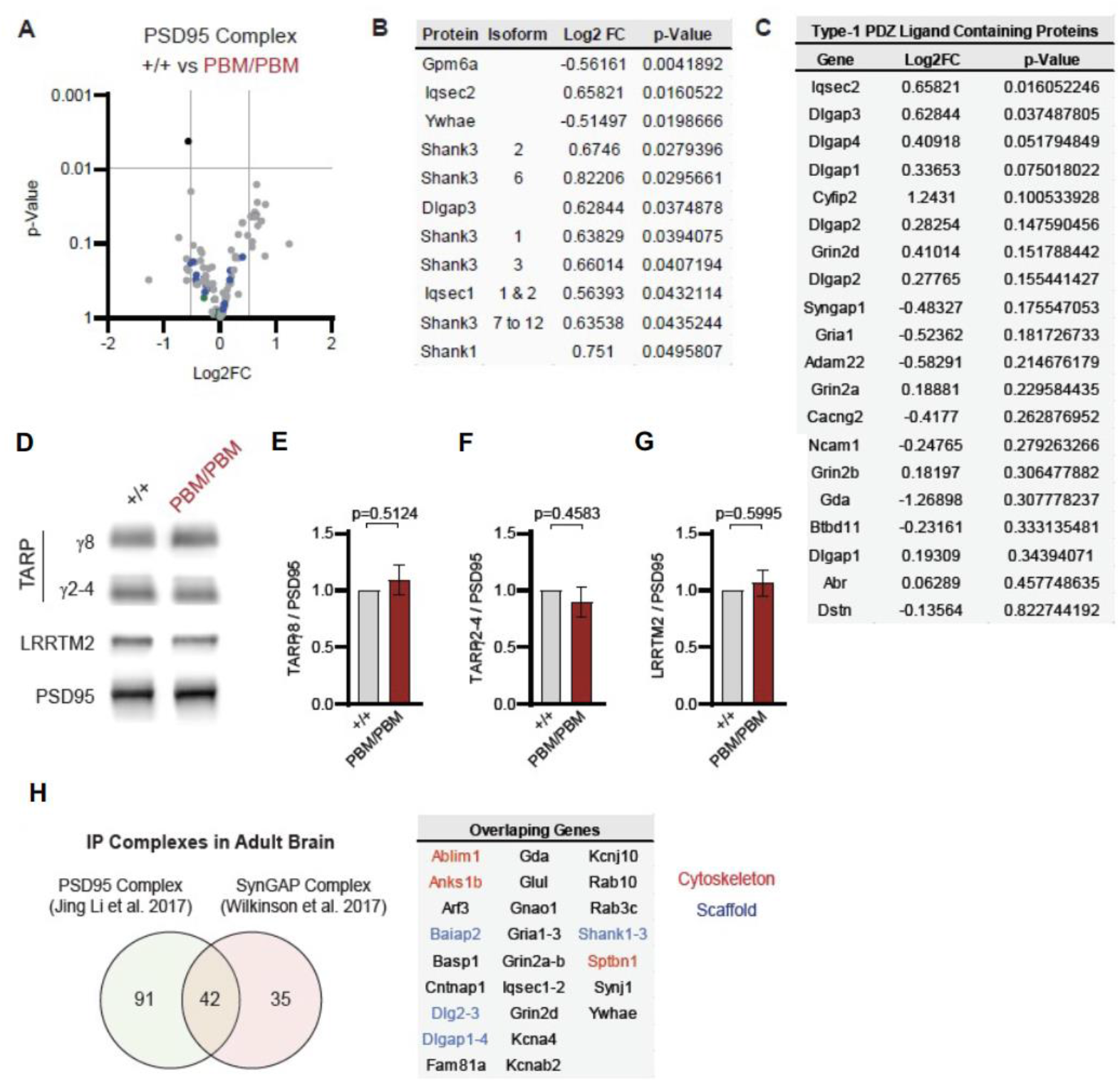
Characterization of native PSD95 complexes from *Syngap1^PBM^* animals. **(A)** Volcano plot demonstrating the label-free quantitative mass-spectrometry profile of the logarithmic difference in protein levels in the immunoprecipitated PSD95 complexes derived from DIV21 +/+ and PBM/PBM cultures in inactivated state. Only Gpm6a (shown in black) was significantly altered beyond p>0.001 cutoff. Blue dots represent proteins with type 1 PDZ-ligands. Green dots represent DLG family proteins. P values were calculated via t-test for each protein. Samples were derived from individual cultures (4 per genotype) which are immunoprecipitated separately. Log2FC was calculated as ratio of PBM/PBM over +/+. **(B)** List of proteins that are differentially expressed beyond p>0.05 cutoff. Note that Iqseq2 and Dlgap3 are PDZ-binding proteins. **(C)** Mass-spectrometry profile of type-1 PDZ binding motif containing proteins in immunoprecipitated PSD95 complex in +/+ vs PBM/PBM inactivated cultures. **(D)** Western blots showing relative expression of TARPs and Lrrtm2 in PSD fractions from adult hippocampi in +/+ vs PBM/PBM. **(E-G)** Quantifications of (D). (E) TARPg8 t(6)=0.6961, p=0.5124. (F) TARPg2-4 t(6)=0.7924, p=0.4583 (G) Lrrtm2 t(6)=0.5542, p=0.5995. Each sample represents hippocampi pooled from 2 mice. **(H)** Comparison of PSD95 and SynGAP IP complexes as reported by (Li et al. 2017 and Wilkinson et al. 2017). Note that PSD95 and SynGAP complexes share diverse range of components involving cytoskeletal and scaffolding proteins.

Given that altering the SynGAP PBM disrupts signaling through NMDARs, we hypothesized that hippocampal CA1 LTP would be disrupted in *Syngap1^PBM/PBM^* mice. The within-train facilitation of responses across the seven theta bursts used to induce LTP did not differ between genotypes **(Fig. 8A)**, indicating that standard measures of induction, including NMDAR channel activation, were not impacted by PBM mutations. However, short-term plasticity (STP; **Fig. 8C, D)** and LTP **(Fig. 8B, E)** were both reduced in *Syngap1^PBM/PBM^* mice. The ratio of LTP/STP was no different between genotypes **(Fig. 8F)**. Blocking NMDAR channel function is known to disrupt both STP and LTP [38]. However, a key measure of NMDA channel function was normal in PBM mutant mice **(Fig. 8A)**. Thus, these data are consistent with the idea that disrupting SynGAP-PDZ binding impairs signaling normally induced downstream of synaptic NMDAR activation. Synaptic plasticity, such as LTP, is thought to contribute importantly to multiple forms of learning and memory. As such, we next measured performance of WT and *Syngap1^PBM/PBM^* mice in a variety of learning and memory paradigms that have previously shown sensitivity in *Syngap1* mouse models, including IRES-TD and β* lines. Behavioral analysis in this line revealed a significant increase in horizontal locomotion in the open field test **(Fig. 8G),** a significantly reduced seizure threshold **(Fig. 8H)**, and significantly reduced freezing during retrieval of a remote contextual fear memory **(Fig. 8I).** Moreover, we also observed impaired acquisition during Morris water maze learning **(Fig. 8J).** Together, these behavioral data indicate that the PBM within SynGAP-α1 splice forms is critical for learning and memory, as well as protecting against seizure.

**Figure 8.**
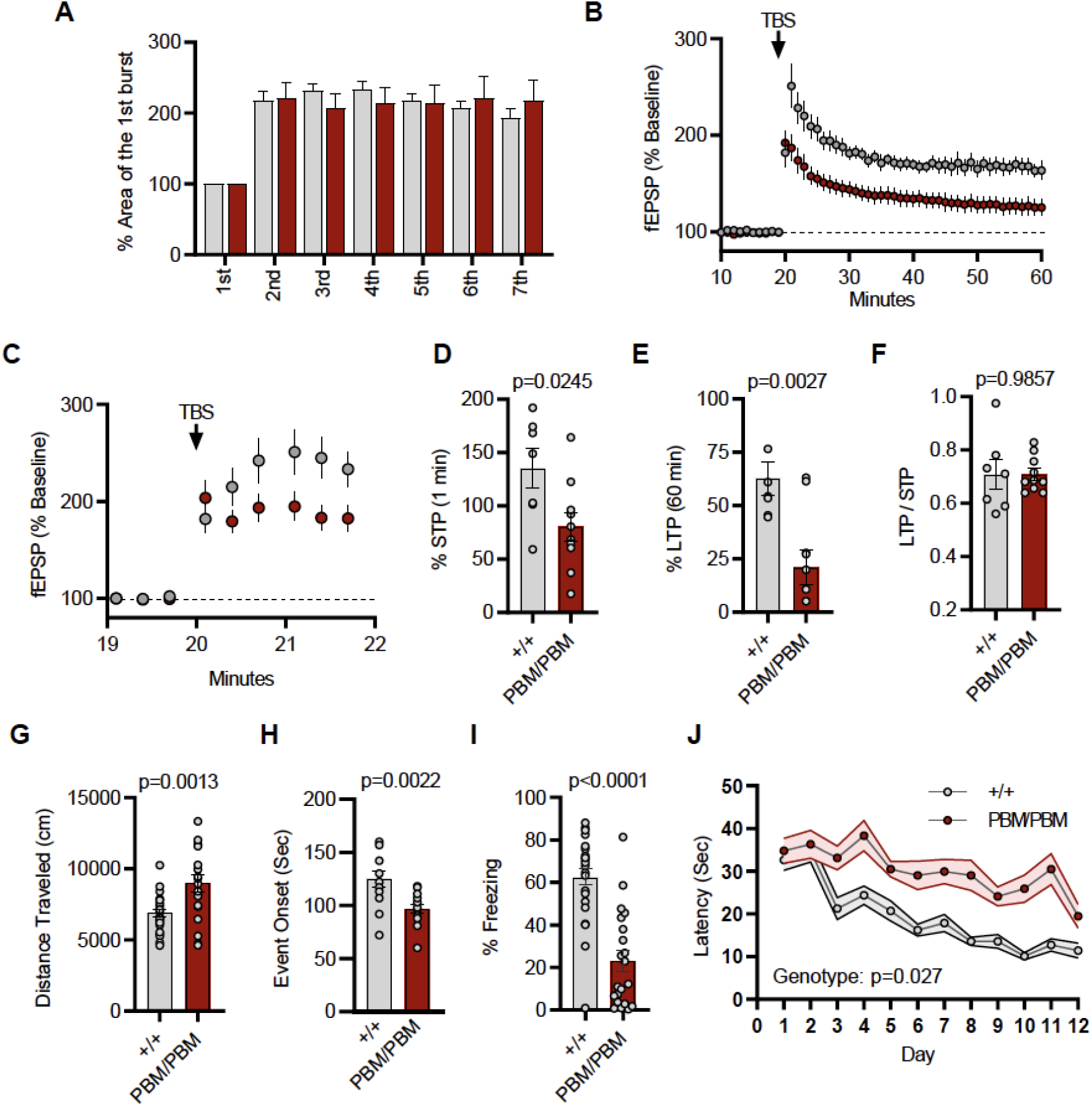
Plasticity and behavior deficits in the *Syngap1^PBM^* mouse line. **A)** Facilitation of burst responses was calculated by expressing the area of the composite fEPSP corresponding to the 2nd theta burst within each train as a fraction of the 1st burst response. No statistically significant difference was found between genotypes. **(B)** Magnitude of long-term potentiation (LTP) following delivery of a single train of five theta bursts. The slope of the fEPSP was normalized to the mean value for a 20 min baseline period; shown are group means and standard errors. The control path, to the same site at which LTP was recorded, received 3/min pulses throughout the session. **(C)** Percent fEPSP during and immediately after the LTP induction. Note that homozygous mutants reach to peak potential immediately following TBS. **(D)** Bar graph shows % potentiation in 1 min after stimulus. t(15)=2.499, p=0.0245 **(E)** Bar graph shows % potentiation in 60 min after stimulus. t(15)=3.594, p=0.0027 **(F)** LTP to STP ratio of individual slices. Note that the level of LTP is proportional to the degree of acute potentiation (1min after stimulus). t(15)=0.01818, p=0.9857. **(G)** Quantification of total distance traveled in OFT. t(45)=3.427, p=0.0013. **(H)** Seizure threshold was measured as the time taken to reach three separate events of 1st clonus (event onset) during the procedure. Unpaired t-test t(25)=3.420 p=0.0022. **(I)** Percent freezing in remote contextual fear memory paradigm. % Freezing: t(45)=6.463, p<0.0001. **(J)** Plots demonstrating latency to find platform across days in Morris Water Maze training session. Statistical significance was determined by using linear mixed model for repeated measures. n=14, +/+ vs PBM/PBM, p=0.027

### Alpha1/2 C-Terminal Isoform expression or function predicts changes in excitatory synapse function

Behavioral results from IRES-TD and PBM mice were consistent with each other, and also consistent with a reduction in all SynGAP isoforms occurring in *Syngap1* conventional heterozygous KO mice. These three mouse lines share a common molecular feature – reduced expression or function of SynGAP-α1 isoforms **(Fig. 1F-I)**. Prior studies have shown that exogenously expressed SynGAP-α1 is a negative regulator excitatory synapse function [25, 39]. Thus, we hypothesized that IRES-TD and PBM mouse lines would express elevated excitatory synapse function, while *Syngap1^β*/β*^* mice, which have enhanced α1 expression, would express reduced synapse function. To test this idea, we performed whole-cell voltage clamp recordings in acute somatosensory cortex slices derived from all three of these lines because these neurons have been shown to be sensitive to *Syngap1* heterozygosity in *ex vivo* slice preparations [21]. PBM mice exhibited a modest increase in *m*EPSCs amplitude and a more substantial increase in *m*EPSC frequency, two measures consistent with enhanced postsynaptic function **(Fig. 9A-C)**. We also observed increased excitatory synapse function (both *m*EPSC amplitude and frequency distributions) in IRES-TD mice **(Fig. 9D-F).** The effects on synapse function from L2/3 SSC neurons observed in these two lines are similar to what has been reported previously in *Syngap1^+/-^* mice [21]. In contrast, *Syngap1^β*/β*^* mice, which have significantly elevated α1 expression, expressed reduced mEPSC amplitude and frequency measurements relative to littermate control slices **(Fig. 9G-I)**, a phenotype consistent with *SynGAP*-α1 overexpression in excitatory neurons [25, 39].

**Figure 9.**
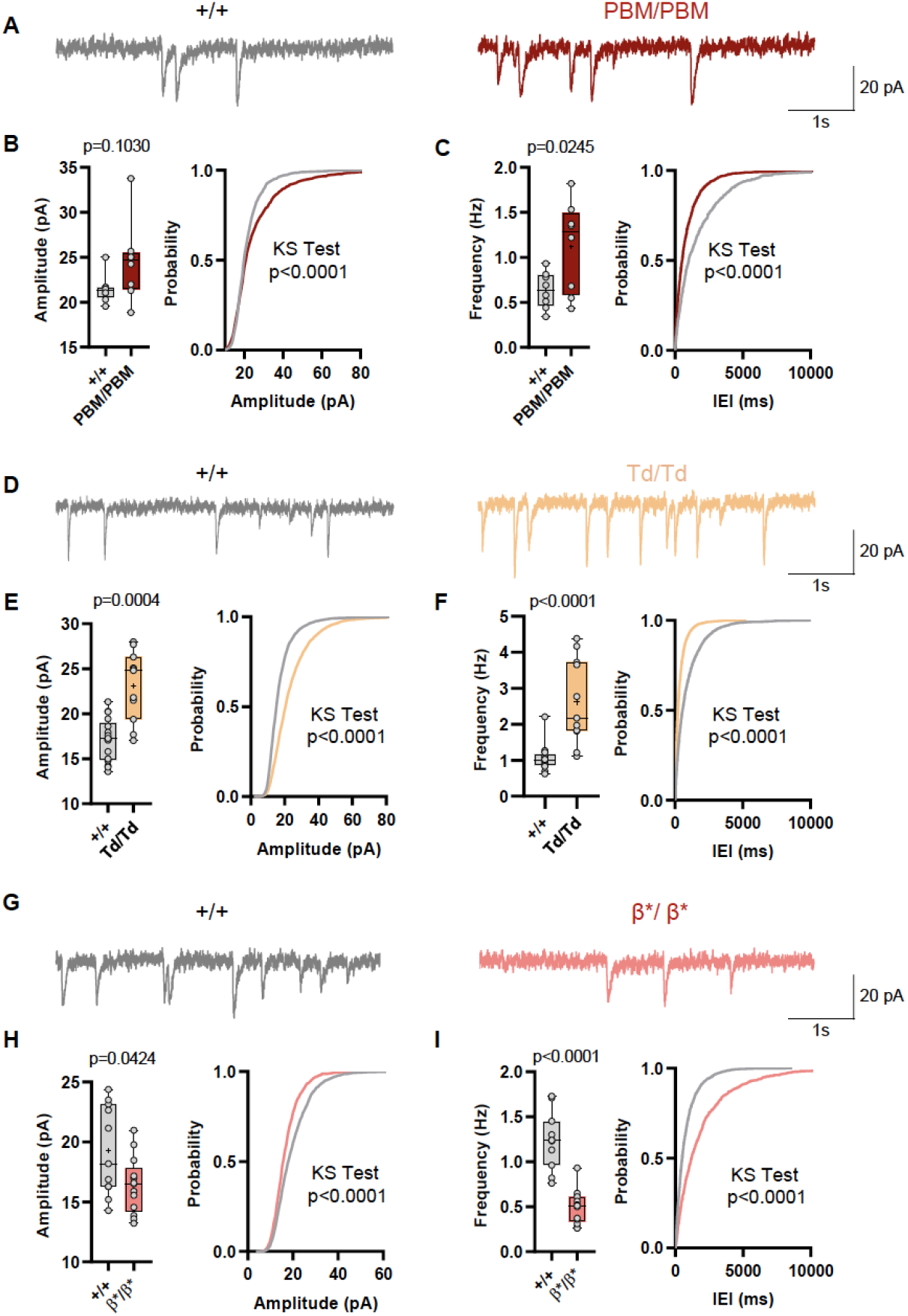
Analysis of excitatory synapse function in *Syngap1^PBM^, Syngap1^β*^*, and *Syngap1^td^* mouse lines. **(A)** Representative mEPSCs traces from L2/3 SSC in +/+ vs PBM/PBM **(B)** Scatter plots and cumulative histograms showing trend towards increase but no significant difference in Amplitudes of mEPSCs +/+ vs PBM/PBM **(C)** Scatter plots and cumulative histograms showing significant increase in frequency of mEPSCs +/+ vs PBM/PBM. Unpaired t test: p=0.0245, n=8 for each genotype. **(D)** Representative mEPSCs traces from L2/3 SSC in +/+ vs Td/Td. **(E)** Scatter plots and cumulative histograms showing significantly increased amplitudes of mEPSCs in +/+ vs Td/Td. Unpaired t test: p=0.0004, n=17 cells for +/+, n=11 cells for Td/Td mice. **(F)** Scatter plots and cumulative histograms showing significant increase in frequency of mEPSCs in +/+ vs Td/Td. Unpaired t test: p<0.0001, n=17 cells for +/+, n=11 cells for Td/Td mice. **(G)** Representative mEPSCs traces from L2/3 SSC in +/+ vs β*/β*. **(H)** Scatter plots and cumulative histograms showing significantly decreased amplitudes of mEPSCs in L2/3 SSC for +/+ vs β*/β*. Unpaired t test: p=0.0424, n=11 cells for +/+, n=13 cells for β*/β*. **(I)** Scatter plots and cumulative histograms showing significant decrease in frequency of mEPSCs in +/+ vs β*/β*. Unpaired t test: p<0.0001, n=11 cells for +/+, n=13 cells for β*/β*.

## Discussion

In this study, we created three distinct mouse lines, each regulating the expression or function of SynGAP protein isoforms **(Fig. 1F-I)**, without appreciable change in total SynGAP expression levels. The overall conclusion from this study is that α-containing SynGAP isoforms promote cognitive functions that support learning/memory, while also protecting against seizure. It is important to understand the relationship between SynGAP isoform function and systems-level manifestations of the different isoforms, such as behavioral expression related to cognitive function and seizure. It has been shown previously that *Syngap1* C-terminal splicing imparts distinct cellular functions of SynGAP proteins [22, 24–26]. Thus, targeting endogenous isoform expression in animal models presents an opportunity to determine to what extent distinct cellular functions of SynGAP could contribute to various intermediate phenotypes present in *Syngap1* mouse models. Given that *SYNGAP1* is a well-established NDD gene and LOF mutations are highly penetrant in the human population [1–3, 5, 6, 8, 40, 41], studying these relationships have the potential to provide much needed insight into the neurobiology underlying human cognitive and behavioral disorders that first manifest during development. Second, there is increasing interest in targeted treatments for patients with *SYNGAP1* disorders due to the penetrance of LOF variants, the relatively homogenous manifestations of the disorder (e.g., cognitive impairment and epilepsy), and the growing number of patients identified with this disorder [42]. Restoring SynGAP protein expression in brain cells is the most logical targeted treatment for this disorder because most known patients have *de novo* variants that cause genetic haploinsufficiency [9]. The most logical therapeutic approach would be to reactivate native expression of the endogenous gene. However, the findings from this study indicate that targeted therapies for *SYNGAP1* disorders that enhance expression of α isoforms may be sufficient to provide a benefit to patients. Indeed, only a modest upregulation of α1/2 expression within a *Syngap1* heterozygous background was sufficient to improve behavioral deficits commonly observed in that mouse line (Fig. 4). Third, the discovery that SynGAP-α1/2 expression/function is pro-cognitive and provides protection from seizure suggests that these isoforms, and the cellular mechanisms that they regulate, could be harnessed to intervene in idiopathic cognitive and excitability disorders, such as neurodegenerative disorders and/or epilepsies with unknown etiology.

Several lines of evidence from this study support the conclusion that SynGAP-α isoform expression or function promotes cognition and seizure protection. IRES-TD and PBM mouse lines each had similar learning/memory and seizure threshold phenotypes, with both mouse lines exhibiting impaired phenotypes related to these two types of behavioral analyses. Indeed, these two mouse lines also shared a common molecular perturbation - reduced expression or function of alpha isoform(s). For example, IRES-TD homozygous mice lacked expression of both α1 and α2 isoforms and these animals exhibited severe phenotypes, including reduced post-weaning survival and dramatically elevated horizontal activity in the open field. Additional phenotypes were also present in heterozygous IRES-TD mice, which underwent more comprehensive testing because of better survival in the post-weaning period. These additional phenotypes included reduced seizure threshold and impaired freezing during a remote contextual fear expression test. PBM homozygous mice had normal expression of SynGAP protein, but lacked a functional domain present exclusively in α1 isoforms, a type-1 PDZ binding domain. PBM homozygous mice shared phenotypes with IRES-TD mice, including impaired remote contextual fear expression, elevated horizontal activity in the open field, and a reduced seizure threshold. These mice also expressed impaired learning during Morris water maze acquisition. Importantly, these behavioral phenotypes are well established in *Syngap1* heterozygous mice [16, 18, 20, 32, 33], indicating that SynGAP protein loss-of-function underlies these abnormalities. Thus, it reasonable to speculate that α isoform LOF is one potential mechanism underlying these behavioral abnormalities. Dysregulation of excitatory synapse function in cortical circuits is one of many possible cellular mechanisms underlying common phenotypes in IRES-TD and PBM mutant mice lines. Whole cell electrophysiology experiments from developing cortical neurons *in situ* from each line revealed evidence of elevated excitatory synapse strength during the known *Syngap1* mouse critical period. Indeed, elevated excitatory synapse strength in developing forebrain glutamatergic neurons is a major cellular outcome present in *Syngap1* heterozygous knockout mice [16, 18, 19, 21]. Moreover, elevated excitatory synapse strength is consistent with impaired cognitive function and reduced seizure threshold.

Studies in the *Syngap1* β* line also support this interpretation. These mice were devoid of SynGAP-β protein expression, yet we did not observe cellular or behavioral phenotypes consistent with *Syngap1* heterozygosity. Rather surprisingly, mice lacking SynGAP-β expression had intermediate phenotypes that opposed what was commonly observed in *Syngap1* heterozygous KO mice (and shared by IRES-TD/PBM lines). For example, β* mice exhibited improved spatial learning in the Morris water maze, reduced horizontal activity in the open field, and an elevated seizure threshold (evidence of seizure protection). These phenotypes were modest in effect size, but highly significant. These phenotypes were reproducible because open field and seizure phenotypes were also present in a separate series of experiments performed in the *Syngap1* heterozygous background. This demonstrates that the impact of the β* allele is penetrant even when expression of isoforms is reduced by half compared to WT mice. As a result, the β* allele partially rescued open field and seizure phenotypes present in *Syngap1^+/-^* mice. For impaired β expression to drive phenotypes, expression of this isoform would be anticorrelated with cognitive function and seizure protection. Put another way, reduced β expression would need to enhance phenotypes and increased expression of these isoforms would need to disrupt them. This outcome is unlikely given that it is inconsistent with phenotypes observed in *Syngap1^+/-^* mice, which have reduced expression of all isoforms, including SynGAP-β.

Phenotypes in β* mice are likely driven by significantly elevated SynGAP-α expression rather than reduced SynGAP-β. Electrophysiological studies in these mice revealed reduced excitatory neuron synaptic strength, a finding consistent with exogenously elevated SynGAP-α1 expression [25, 39]. Moreover, these synapse-level results are consistent with seizure protection observed in β* mice. Phenotypes in PBM mice also support this hypothesis. This model does not have altered t-SynGAP expression, or a change in β expression. Yet, the behavioral- and synapse-level phenotypes are consistent with those observed in IRES-TD and *Syngap1^+/-^* mice. The observation that α isoforms promote cognitive function and seizure protection are consistent with known molecular functions of these isoforms, at least with respect to regulation of synapse strength and resultant impacts on neural circuit function. For example, α1 imparts SynGAP with the ability to undergo liquid-liquid phase transitions [34]. This biophysical process is associated with regulation of Ras signaling in dendritic spines required for AMPA receptor trafficking that supports use-dependent synapse plasticity [17, 22]. Input-specific plasticity is crucial during development to sculpt the assembly of neural circuits [43], while also being important in mature circuits to promote experience-dependent changes in already-established circuitry [44].

A consensus is emerging that baseline synaptic phenotypes related to *Syngap1* gene expression are dominated by the ability of both α1 and α2 isoforms to suppress excitatory synapse function. Studies from several research groups have shown that SynGAP-α1 is a negative regulator of excitatory synapse structure and function [17, 22, 25, 26, 39]. In contrast, the role of α2 isoform protein function on excitatory synapse structure/function is less clear. One study suggested that α2 has an opposing function relative to α1 within excitatory synapses, with the former acting as an enhancer, rather than a suppresser, of excitatory synapse function [24]. However, a more recent study demonstrated that α2 has a similar, albeit less robust ability to suppress AMPA receptor content within dendritic spines [22], indicating that it too can act as a negative regulator of synapse function. Our results here support the view that both α1 and α2 can act as suppressors of excitatory synapse function. In our studies, α1 and α2 were both co-regulated in the IRES-TD and β* lines, with both isoforms downregulated in the former and upregulated in the latter. In both mouse lines, baseline excitatory synapse strength was inversely proportional to expression levels of α1/2 isoforms. If α1 and α2 had opposing functions at the synapse level, then co-regulation of both isoforms would be expected to lead to no significant differences in synapse function.

It is important to note that our interpretation that β* mouse phenotypes are most likely driven by changes in α isoforms does not preclude a fundamental role of β in sculpting neural systems, or that reduced expression of this isoform in *Syngap1^+/-^* mice has no role in disease pathobiology. Rather, our results highlight the importance of endogenous α isoforms in regulating excitatory synapse function and associated behavioral outcomes. What is known about the function of other C-terminal protein variants, such as β and γ? A recent study suggested that β and γ isoforms lack the ability to regulate excitatory synapse function, further strengthening the idea that α isoforms account for *Syngap1*-dependent regulation of excitatory synapse function [22]. However, *Syngap1* is known to regulate additional cellular process beyond regulation of excitatory synapse function, such as dendritic morphogenesis and patterning *in vivo* [18, 20, 21]. Evidence suggests that all isoforms can regulate dendritic morphogenesis *in vitro*, though SynGAP*-*β was shown to be a stronger regulator of this process relative to the other C-terminal isoforms [22]. *In vivo*, β was found to be expressed earlier in development and to be less enriched in the postsynaptic density compared to other variants [23]. Thus, β is well positioned to regulate non-synapse related neuronal processes. Future studies will be required to elucidate the specific cellular functions of non-alpha isoforms and how they contribute to the development of neural function and behavior. Given the complexities of *Syngap1* regulation on dendritic morphogenesis [20, 21], and the direct linkage between dendritic morphogenesis and circuit function in cortex in *Syngap1* mutant animals [21], future studies on the function of individual isoforms would ideally be carried out *in vivo* in developing animals.

## Acknowledgements

This work was supported in part by NIH grants from the National Institute of Mental Health (MH096847 and MH108408 to G.R., MH115005 and MH113949 to M.P.C, and MH105400 to C.A.M.), the National Institute for Neurological Disorders and Stroke (NS064079 to G.R.), the Eunice Kennedy Shriver National Institute of Child Health and Human Development (HD089491 to G.L.), the National Institute for Drug Abuse (DA034116 and DA036376 to C.A.M.), the Spanish Ministerio de Ciencia, Innovación y Universidades (BFU2012-34398, BFU2015-69717-P, RTI2018-097037-B-100, RYC-2011-08391p and IEDI-2017-00822) and the Catalan Government (AGAUR SGR14-297 and 2017SGR1776). M.K. was supported by Autism Speaks Weatherstone Pre-Doctoral fellowship (10646). G.G. was supported by a predoctoral fellowship from the Spanish Ministerio de Educación (BES-2013-063720). V.A. was supported by a training Fellowship from Leon and Friends.

## Author Contributions

M.K. performed experiments, designed experiments, analyzed data, co-wrote the manuscript, and edited the manuscript. V.A performed experiments, designed experiments, analyzed data, and edited the manuscript. T.K.C. performed experiments, designed experiments, analyzed data, and edited the manuscript. C.R. performed experiments, designed experiments and analyzed data. A.A.L performed experiments, designed experiments, and analyzed data. J.L designed experiments and analyzed data. B.W. performed experiments, designed experiments, and analyzed data. N.H. performed experiments and designed experiments. N.G. performed experiments and designed experiments. A.R. analyzed data. G.G. performed experiments. Y.A. performed experiments. A.B. designed experiments, analyzed data, and interpreted data. M.P.K. designed experiments, analyzed data, and interpreted data. G.L. designed experiments, analyzed data, and interpreted data. C.A.M. designed experiments, interpreted data, and edited the manuscript. G.R. conceived the study, designed experiments, interpreted data, co-wrote the manuscript, and edited the manuscript.

## Declaration of Interests

The authors declare no competing financial interests.

## Materials & Methods

### Animals

This study was performed in strict accordance with the recommendations in the Guide for the Care and Use of Laboratory Animals of the National Institutes of Health. All of the animals were handled according to approved institutional animal care and use committee (IACUC) protocols of The Scripps Research Institute.

*Syngap1*^PBM^ and *Syngap1^Td^* mice were constructed in collaboration with genOway (France). The targeting vector was electroporated into ES cells derived from the inner cell mass of 3.5 days old C57BL/6N embryos. Cells were then subjected to negative and/or positive selection(s) before the presence of the correct recombination event was validated by PCR and Southern blot. ES cell clones with verified mutations were injected into blastocysts which were implanted into pseudo-pregnant females to obtain chimeras. Chimeric mice were bred with C57BL/6 Cre-deleter mice to excise the Neomycin selection cassette and to generate heterozygous mice carrying the Neo-excised knock-in allele. Progeny were genotyped by PCR. The recombinase-mediated excision event was further validated by Southern blot using 5’ external probes. Knock-in lines were maintained on C57BL/6J background and bred for 3 generations prior to experimental use. *Syngap1* ^PBM^ animals were genotyped using the following primers, which amplified the locus spanning the LoxP site: Fwd: 5’-ctggttcaaaggctcctggta-3’ Rev: 5’-ctgtttgtttctcacctccaggaa-3’. This combination yielded a 61bp product in WT and 120bp product in knock-in alleles. *Syngap1^Td^* line were genotyped using the primers amplifying the locus including the TdTomato cassette: Fwd: 5’-AGATCCACCAGGCCCTGAA-3’ Rev: 5’-GTCTTGAACTCCACCAGGTAGTG-3’

*Syngap1-β\** mice were constructed in collaboration with the Scripps Research Genetics core facility. To selectively disrupt SynGAP-β expression, exon19a splice acceptor site “AAG” was mutated into “ACG”. To introduce the point mutation, purified CRISPR/Cas9 protein combined with gRNA and donor DNA was injected to ~100 zygotes and implanted into surrogate mice. A 200 bp PAGE purified ss-oligo repair template centering the CRISPR cut site was used as donor DNA. Recombination events were detected by PCR and Sanger sequencing of the DNA isolated from tails of F0 potential founders. This process identified 2 chimeric mice with evidence of the targeted nucleotide variants. Chimeras were then bred with C57BL6/J and resultant heterozygous F1 mice were used to start the colony. Because CRISPR carries a risk of off-target genomic effects, prior to any downstream experiments, this line was further crossed into C57BL6/J for >3 generations.

### Transcriptomics

PND7 mice forebrains (Cortex + hippocampus) were immediately removed and stored in RNALater (Thermo, AM7020). mRNA was isolated with RNeasy mini kit (74104, Qiagen). RNA integrity was measured using Agilent 2100 Bioanalyzer (RIN value >= 9.2 for each sample). Library preparation and sequencing on the Illumina NextSeq 500 were performed by the Scripps Florida Genomics Core. De-multiplexed and quality filtered raw reads (fastq) were trimmed (adaptor sequences) using Flexbar 2.4 and aligned to the reference genome using TopHat version 2.0.9 (Trapnell et al., 2009). HT seqcount version 0.6.1 was used to generate gene counts and differential gene expression analysis was performed using Deseq2 (Anders and Huber, 2010). DeSeq2 identified differentially expressed genes (DEGs) with a cutoff of 1.5 fold change and an adjusted p-value of less than 0.05 (Love et al., 2014). Paired end reads mapped to the first 30 bases of Exon21 was used to determine the ratio of Exon21a (results in SynGAP-α2) vs Exon21b (results in SynGAP-α1) splicing events.

### Cell Culture

#### Cell lines

HeLa Cells (Kind gift of Michael Farzan) and HEK293T Cells (Kind gift of Joseph Kissil) were cultured in DMEM media containing 10% fetal bovine serum and penicillin/streptomycin.

#### Primary forebrain cultures

Dissociated forebrain cultures were prepared from newborn WT and homozygous littermates of the PBM line as previously described (Bedouin 2012). Briefly, forebrains were isolated and incubated with a digestion solution containing papain for 25 min at 37 °C. Tissues were washed and triturated in Neurobasal medium containing 5% FBS. Cells were plated on poly-D-lysine at a density of 1,000 cells per mm^2^. Cultures were maintained in Neurobasal A media (Invitrogen) supplemented with B-27 (Invitrogen) and Glutamax (Invitrogen). At DIV4 cells were treated with FuDR to prevent glial expansion. The cells were sparsely labeled by administration of AAVs (CamKII.Cre, 10^4^vg/ml, Addgene # 105558-AAV9 and CAG.Flex.EGFP, 10^8^vg/ml, Addgene #28304-PHPeB) at DIV 9-10 and processed for experiments 10-11 days later.

### *In situ* Colocalization Assay

HeLa cells were plated on glass coverslips and transfected with PSD95-tRFP (Plasmid #52671, Addgene) and/or EGFP-tagged SynGAP C-terminal constructs (EGFP-CCα1 or EGFP-CCPBM plasmids (made in house) were co-transfected into HeLa cells using lipofectamine 2000 according to manufacturer instructions. Cells were then fixed with 4% PFA and washed multiple times with PBS prior to mounting with Prolong Gold with DAPI (P36931, Thermo). Confocal stacks spanning entire cells were obtained using UPlanSApo 100× 1.4 NA oil-immersion objective mounted on Olympus FV1000 laser-scanning confocal microscope using Nyquist criteria for digital imaging. Maximum intensity projections were used for the analysis. Nuclei of cells were defined by DAPI staining, and the EGFP-CC nuclear localization was calculated as the EGFP (colocalized with nucleus) / EGFP (within entire cell perimeter).

### PSD95-SynGAP Co-IP Assay

PSD95-tRFP (Plasmid #52671, Addgene) and/or full length EGFP-SynGAPα1/PBM (made in house) plasmids were transfected in HEK293T cells using Lipofectamine 2000. Cells were homogenized with Pierce IP Lysis buffer (87787, Thermo) containing protease & phosphatase inhibitors. Lysates were then incubated for 2hrs at RT with 1.5mg Dynabeads (10004D, Thermo) functionalized with 10ug of anti-PSD95 (Thermo, MA1-045) or IgG control (ab18415, Abcam). After extensive washing, immunoprecipitated proteins were eluted with Leammeli buffer at 70C for 10min with agitation. Eluted proteins were detected via western blot using PSD-95 (Thermo, MA1-045) and SynGAP (D20C7, CST) antibodies.10% of the input and 20% of IP elute were used for each sample.

### *In Vitro* Treatments

To silence neuronal activity and block NMDAR signaling, cultures were treated for 3hrs with 1 μM TTX and 200 μM APV. To induce chemical LTP, Cells were thoroughly washed and perfused with basal ECS (143 mM NaCl, 5 mM KCl, 10 mM HEPES (pH 7.42), 10 mM Glucose, 2 mM CaCl_2_, 1 mM MgCl_2_, 0.5 μM TTX, 1 μM Strychnine, and 20 μM Bicuculline) for 10 min. Then magnesium free ECS containing 200 μM Glycine (or 10 μM Glycine for weak cLTP) was applied for 10 min. Cells were then washed with and incubated in basal ECS for additional 10 min prior to downstream application.

### Subcellular Fractionation

#### From tissue

Frozen hippocampi or cortex were homogenized using a Teflon-glass homogenizer in ice-cold isotonic solution (320 mM sucrose, 50 mM Tris pH 7.4, phosphatase & protease inhibitors). The homogenate was then centrifuged at 1,000g for 10min at 4 °C. The supernatant (S1) was centrifuged at 21,000g for 30min. The pellet (P2) was resuspended in isotonic buffer and layered on top of a discontinuous sucrose density gradient (0.8M, 1.0M or 1.2M sucrose in 50mM Tris pH 7.4, +inhibitors) and centrifuged at 82,500g for 2hr at 4°C. The interface of 1.0M and 1.2M sucrose was collected as a synaptosomal fraction. Synaptosomes were diluted using 50mM Tris pH7.4 (+inhibitors) to bring the sucrose concentration to 320mM. The diluted synaptosomes were then pelleted by centrifugation at 21000g for 30min at 4°C. The synaptosome pellet was then resuspended in 50mM Tris pH 7.4 and then mixed with an equal part 2% Triton-X (+inhibitors). This mixture was incubated at 4 °C with rotation for 10min followed by centrifugation at 21,000xg for 20min to obtain a supernatant (Syn/Tx) and a pellet (PSD).

#### From primary culture

Cultured neurons (DIV 18-21), were homogenized by passage through 22G needle 10 times in ice-cold isotonic buffer (320 mM sucrose, 50 mM Tris, protease & phosphatase inhibitor mix). Homogenates were centrifuged at 1,000 × *g* for 10 min at 4 °C. The supernatant (S1) was centrifuged at 15,000 × *g* for 20 min at 4 °C to obtain the crude membrane (P2 fraction). The P2 pellet was resuspended with ice-cold hypotonic buffer (50 mM Tris, protease & phosphatase inhibitor mix) and was incubated for 30 min at 4C. Then the sample was centrifuged 21,000 x g for 30min to obtain synaptic plasma membrane (SPM) fraction. SPM is reconstituted in hypotonic buffer then equal volume of hypotonic buffer with 2% Triton-X was added and the mixture was incubated 15min on ice. Lysates were centrifuged at 21,000*g* for 30 min at 4 °C to obtain a soluble fraction (Syn/Tx) and a pellet (PSD), which was resuspended in 50 mM Tris containing 0.5% SDS. To completely solubilize PSD fraction, we’ve briefly sonicated and heated samples to 95 °C for 5min.

### Immunoblotting

Protein lysates were extracted from the hippocampi or cortices of adult mice and dissected in ice-cold PBS containing Phosphatase Inhibitor Cocktails 2 and 3 (Sigma-Aldrich, St. Louis, MO) and Mini-Complete Protease Inhibitor Cocktail (Roche Diagnostics) and immediately homogenized in RIPA buffer (Cell Signaling Technology, Danvers, MA), and stored at −80 °C. Sample protein concentrations were measured (Pierce BCA Protein Assay Kit, Thermo Scientific, Rockford, IL), and volumes were adjusted to normalize microgram per microliter protein content. For phospho-protein analysis, *in vitro* cultures were directly lysed with laemmeli sample buffer, sonicated and centrifuged to minimize DNA contamination. 10 μg of protein per sample were loaded and separated by SDS-PAGE on 4-15 % gradient stain-free tris-glycine gels (Mini Protean TGX, BioRad, Hercules, CA), transferred to low fluorescence PVDF membranes (45 μm) with the Trans-Blot Turbo System (BioRad). Membranes were blocked with 5% powdered milk (BSA for phospho-proteins) in TBST and probed overnight at 4 °C with the following primary antibodies: Pan-SynGAP (Thermo, PA1-046), SynGAP-α1 (Millipore, 06-900), SynGAP-α2 (abcam, ab77235), SynGAP-β (Kind gift of Rick Huganir), PSD-95 (Thermo, MA1-045), Synaptophysin (Novus, NB300-653), pERK (CST, 9106), ERK (CST, 4696), GluA1 (Millipore, MAB2263), phospho-serine845 GluA1 (Millipore, AB5847), TARP (Millipore, Ab9876), LRRTM2 (Thermo Pierce, PA521097).

### Immunocytochemistry

*For SynGAP – PSD95 colocalization*, neurons were fixed in 4% PFA, 4% sucrose for 5 min at RT and treated with MetOH for 15min at −20°C. The cells were then washed with PBS and permeabilized in PBS 0.2% TritonX-100 for 10 min. Samples were then blocked for 1 hr and probed for SynGAP (D20C7, CST) and PSD95 (MA1-045, Abcam) overnight. After PBS washes, samples were probed with appropriate secondary antibodies for 1 hr in the dark at room temperature. The coverslips were then washed, mounted (Prolong Glass) and cured. Confocal stacks were obtained. For analysis, maximum intensity Z projection was obtained from each confocal image. Individual synapses were traced as PSD95 positive puncta selected using an arbitrary threshold which was kept constant across all images. Mean SynGAP and PSD95 signals were measured from individual synapses. *For surface GluA1 staining*, neurons were immediately fixed in ice-cold pH 7.2 4% PFA, 4% sucrose for 20 min on ice. Then, samples were washed three times with ice-cold PBS and blocked for 1 hr min in PBS containing 10% NGS. Cells were then incubated overnight with a primary antibody targeting the extracellular N terminus of GluA1 (MAB2263, Millipore) and then washed with 10% goat serum twice to remove excess primary antibody. After PBS washes, Alexa dye–conjugated secondary antibodies were added for 1 hr in the dark at room temperature. The coverslips were then washed, mounted (Prolong Glass) and cured. Surface GluA1 levels were measured from manually traced individual dendritic spines from maximum intensity Z projection images using EGFP channel (cell fill). All confocal stacks were obtained for 6–12 individual fields from multiple coverslips per culture with UPlanSApo 100× 1.4 NA oil-immersion objective mounted on Olympus FV1000 laser-scanning confocal microscope using Nyquist criteria for digital imaging. 40-80 μm stretches of secondary dendrites in neurons with pyramidal morphology were imaged.

### PSD95 Immunoprecipitation and Mass Spectrometry

Harvested neurons were lysed in DOC lysis buffer (50 mM Tris (pH 9), 30 mM NaF, 5 mM sodium orthovanadate, 20 mM β-glycerol phosphate, 20 μM ZnCl_2_, Roche complete, and 1% sodium deoxycholate). The lysate was then centrifuged at 35,000 RPM for 30 minutes at 4°C and lysate containing 1 mg of protein was incubated with 2 μg Psd95 antibody (Neuromab, catalog # 75-048) at 4°C overnight with rotation. The following day, IPs were incubated with Dynabeads protein G (Thermo Fisher Scientific, catalog # 10004D) for 2 hours at 4 degrees Celsius. IPs were then washed three times with IP wash buffer (25 mM Tris (pH 7.4), 150 mM NaCl, 1 mM EDTA, and 1% Triton X-100). Dynabeads were re-suspended in 2X LDS sample buffer and incubated at 95 degrees Celsius for 15 minutes for elution. The eluate was incubated with DTT at a final concentration of 1 mM at 56°C for 1 hour followed by a 45-minute room temperature incubation with Iodoacetamide at a final concentration of 20 mM.

Samples were loaded onto 4 – 12% Bis-Tris gels and separated at 135V for 1.5 hours. Gels were stained with InstantBlue (Expedeon, catalog # 1SB1L) to visualize bands. The heavy and light chains of Immunoglobulin were manually removed. Gels were then destained using 25% ethanol overnight. Gel lanes were cut, individual gel slices were placed into 96 well plates for destaining, and peptide digestion was completed at 37 degrees Celsius overnight. Peptides were extracted with acetonitrile, dried down, and then desalted using stage tips. All LC-MS experiments were performed on a nanoscale UHPLC system (EASY-nLC1200, Thermo Scientific) connected to an Q Exactive Plus hybrid quadrupole-Orbitrap mass spectrometer equipped with a nanoelectrospray source (Thermo Scientific). Samples were resuspended in 10uL of Buffer A (0.1% FA) and 2uL were injected. Peptides were separated by a reversed-phase analytical column (PepMap RSLC C18, 2 μm, 100 Å, 75 μm X 25 cm) (Thermo Scientific). Flow rate was set to 300 nl/min at a gradient starting with 3% buffer B (0.1% FA, 80% acetonitrile) to 38% B in 110 minutes, then ramped to 75% B in 1 minute, then ramped to 85% B over 10 minutes and held at 85%B for 9 minutes. Peptides separated by the column were ionized at 2.0 kV in the positive ion mode. MS1 survey scans for DDA were acquired at resolution of 70k from 350 to 1,800 m/z, with maximum injection time of 100 ms and AGC target of 1e6. MS/MS fragmentation of the 10 most abundant ions were analyzed at a resolution of 17.5k, AGC target 5e4, maximum injection time 65 ms, and an NCE of 26. Dynamic exclusion was set to 30 s and ions with charge 1 and >6 were excluded. The maximum pressure was set to 1,180 bar and column temperature was constant at 50°C. Proteome Discoverer 2.2 (Thermo Fisher Scientific) was used to process MS data and analyzed using Sequest HT against Uniprot mouse databases combined with its decoy database. With respect to analysis settings, the mass tolerance was set 10 parts per million for precursor ions and 0.02 daltons for fragment ions, no more than two missed cleavage sites were allowed, static modification was set as cysteine carbamidomethylation, and oxidation of methionine was set as a dynamic modification. False discovery rates (FDRs) were automatically calculated by the Percolator node of Proteome Discoverer with a peptide and protein FDR cutoff of 0.01. Label free quantification was performed using Minora node in Proteome Discoverer. Abundances of identified PSD95 interacting proteins in WT and mutant neurons were compared using relative abundances such that proteins with a fold change in abundance ratio of > 2.0 or < 0.5 were considered to be differentially associated to PSD95.

### Hippocampal LTP and Extracellular Recordings

Acute transverse hippocampal slices (350 μm) were prepared using a Leica Vibroslicer (VT 1000S), as described previously (Babayan et al., 2012). Slices were cut into ice cold, choline chloride artificial cerebral spinal fluid (ACSF) containing (in mM) 110 choline chloride, 2.5 KCl, 1.25 NaH2PO4, 5 MgSO4, 25 NaHCO2, 25 glucose, 11.6 ascorbic acid, and 3.1 pyruvic acid and rinsed at room temperature for ~3 min in a high magnesium aCSF solution containing: 124 NaCl, 3 KCl, 1.25 KH2PO4, 5 MgSO4, 26 NaHCO3, and 10 dextrose. Slices were then transferred to an interface recording chamber maintained at 31±1°C, oxygenated in 95% O2/ 5% CO2 and constantly perfused (60-80 ml/h) with normal ACSF (in mM; 124 NaCl, 3 KCl, 1.25 KH2PO4, 1.5 MgSO4, 2.5 CaCl2, 26 NaHCO3, and 10 dextrose). Slices equilibrated in the chamber for approximately 2 hours before experimental use. Field excitatory postsynaptic potentials (fEPSPs) were recorded from CA1b stratum radiatum using a single glass pipette (2-3 MΩ). Bipolar stainless-steel stimulation electrodes (25 μm diameter, FHC) were positioned at two sites (CA1a and CA1c) in the apical Schaffer collateral-commissural projections to provide activation of separate converging pathways of CA1b pyramidal cells. Pulses were administered in an alternating fashion to the two electrodes at 0.05 Hz using a current that elicited a 50% maximal response. After establishing a 10-20 min stable baseline, long-term potentiation (LTP) was induced in the experimental pathway by delivering 7 ‘theta’ bursts, with each burst consisting of four pulses at 100 Hz and the bursts themselves separated by 200 msec (i.e., theta burst stimulation or TBS). The stimulation intensity was not increased during TBS. The control pathway received baseline stimulation (0.05Hz) to monitor the health of the slice. The fEPSP slope was measured at 10–90% fall of the slope and all values pre- and post-TBS normalized to mean values for the last 10 min of baseline recording. Baseline measures for all groups included paired-pulse facilitation and input/output curves.

### *Ex vivo* whole-cell electrophysiology

Acute coronal slices (350 μm) were prepared from 10-14 days old mice for 3 mouse lines. Ice-cold cutting solution was used for slice preparation and contained the following (in mM): 119 NaCl, 2.5 KCl, 1.3 MgSO4, 2.5 CaCl2, 1 NaH2PO4, 11 D-glucose and 26.3 NaHCO3, pH 7.4, 300-310 mOsm bubbled with 95%CO2 and 5%O2. The slices were then warmed to 37°C for an hour approximately in standard artificial cerebrospinal fluid (aCSF), composed of (mM): 125 NaCl, 2.5 KCl, 24 NaHCO3, 2 CaCl2, 1.25 NaH2PO4, 2 MgSO4, and 10 D-Glucose, and equilibrated with 95 % O2 and 5 % CO2 (pH 7.4, ~300 mOsm). Following this, slices were maintained in bubbled aCSF at room temperature until transferred to a submerged-type recording chamber (Warner Instruments, Hamden, CT). All experiments were performed at 32°C±2 (perfusion rate of 2-3 mL/min). Whole-cell patch clamp experiments were conducted from visually identified L2/3 neurons using infrared DIC optics. L2/3 excitatory cells were identified by their soma shape and their location ~ 150 uM below the L1-L2 boundary. Regular spiking was confirmed in current clamp and miniature excitatory postsynaptic current (mEPSC) were recorded from identified cells for 5 sweeps each lasting a minute, using the following internal solution (in mM): 120 CsCl, 10 K-HEPES, 10 EGTA, 5 QX314-Br, 4 Mg-ATP, 0.3 Na-GTP, 4 MgCl2 (pH 7.3, 290-295 mOsm). Perfusion solution aCSF was supplemented with 100 μM picrotoxin and 1 μM TTX. Cells with access resistance >20 MΩ or were unstable (>20 % change) were discarded from further analysis. Recordings were made using borosilicate glass pipettes (3-6 MΩ; 0.6 mm inner diameter; 1.2 mm outer diameter; Harvard Apparatus). All signals were amplified using Multiclamp 700B (Molecular Devices, Sunnyvale, CA), filtered at 4 KHz, digitized (10 KHz), and stored on a personal computer for off-line analysis. Analog to digital conversion was performed using the Digidata 1440A system (Molecular Devices). Data acquisition and analyses were performed using pClamp 11.2software package (Clampex and Clampfit programs; Molecular Devices) and minianalysis (Synaptosoft). The events were considered mini-EPSCs if the peak of an event was >5 pA.

### Behavior

At weaning, four mice were randomly allocated to one cage with respect to genotype with males and females being housed separately. Randomization of cage allocation was restricted in that, as much as possible, mice from the same litter were placed in different cages so that no single litter was overrepresented in any single experiment. Cages utilized for behaviors contained cardboard pyramidal-shaped huts with two square openings on opposing sides of the hut for the purposes of environmental enrichment and to assist with transfers from home cages to behavioral apparatuses. All mice were handled for several minutes on three consecutive days prior to commencement of behavioral testing. Tails were marked for easy identification and access from home cages during testing. Experimenters were blind to mouse genotype while conducting all tests.

#### Flurothyl-induced seizures

Flurothyl-induced seizure studies were performed based on prior studies with some modifications [16, 18, 45]. Briefly, experiments were conducted in a chemical fume hood. Mice were brought to the experimental area at least 1 h before testing. To elicit seizures, individual mice were placed in a closed 2.4-L Plexiglas chamber and exposed to 99% Bis (2,2,2-triflurothyl) ether (Catalog# 287571, Sigma-Aldrich, St. Louis, MO). The flurothyl compound was infused onto a filter paper pad, suspended at the top of the Plexiglas chamber through a 16G hypodermic needle and tube connected to a 1 ml BD glass syringe fixed to an infusion pump (KD Scientific, Holliston, MA, USA, Model: 780101) at a rate of 0.25 ml/min. The infusion was terminated after the onset of a hind limb extension that usually resulted in death. Cervical dislocation was performed subsequently to ensure death of the animal. Seizure threshold was measured as latency (s) from the beginning of the flurothyl infusion to the beginning of the first myoclonic jerk.

#### Morris water maze

Mice were run in a standard comprehensive Morris water maze paradigm including a cue test with a visual platform and an acquisition protocol with a hidden platform. All phases of the paradigm were run in a dedicated water maze room in the Scripps Florida Mouse Behavior Core. A water maze system including a plastic white opaque pool (Cat# ENV-594M-W, Med Associates), measuring ~122cm diameter at the water surface, supported by a stand (ENV-593M-C) and equipped with a floor insert (ENV-595M-FL) covering a submerged heater was utilized for all water maze experimentation. An adjustable textured platform (17.8 cm diameter, ENV-596M) was placed atop the floor insert in one of two different quadrants, depending on the specific phase of the paradigm (NW quadrant for initial training and probe test and SE quadrant for reversal training and probe tests), for mice to escape the water. Water temperatures were controlled to 22.5 ± 0.5 °C using a built-in heater and monitored with a digital temperature probe. This water temperature motivated the mice to escape the water without eliciting hypothermic conditions. The tank was emptied, cleaned and refilled once every three days to avoid unsafe accumulation of bacteria. Water was made opaque by the addition of a white opaque non-toxic paint (Crayola) forcing mice to utilize extra-maze cues when locating the hidden platform (0.5 cm beneath the surface of the water). These spatial cues (large black cardboard circle, star, square, white X on black background) were placed on the walls of the room at different distances from the pool. The pool edge was demarcated with directional units (W, N, E, S) to aid assignment of invisible platform “quadrants” to the pool arena outlined by the video tracking system. Various strip lights were positioned on the walls near the ceiling to allow for a moderate level of lighting (200 lux), enough for the mice to see the extra-maze cues adequately without eliciting undue anxiety. Thirty minutes prior to commencement of daily trials, the lights and heater were turned on, and mouse home cages were placed on heating pads on a rack in the water maze room to provide a warm place for the mice between trials. Cage nestlets were replaced with strips of paper towels to better facilitate drying after trials. Mice were monitored during trials for signs of distress and swimming competence. None of the mice tested had swimming issues, and floating was discouraged with gentle nudges. Mice received four trials per day during cue and acquisition phases and one trial per day for probe trials. Three cages (12 mice) were run at a time such that ITIs for each day lasted about 20 minutes with trial duration lasting until the mouse found the platform or a maximum of 60 s. Each trial commenced when the mouse was automatically detected in the pool by the tracking system (Ethovision, Noldus). Each mouse was lowered into the pool facing its edge at one of the four directional units (W, N, E, S) in a clockwise manner, with the first of the four trials starting closest to the platform (“NW quadrant”), which was positioned in the central area of the quadrant dictated by the tracking system. This same series of daily trial commencements were followed for all mice for each of the cue tests, acquisition protocol, and reversal protocol. If the mouse did not locate the platform in 60 s, the experimenter’s hand guided them to the platform. Because the mice are eager to escape the water, the mice quickly learned to follow hand direction to the platform, minimizing physical manipulation of the animals during the trials. Mice were allowed 15 seconds on the platform at the end of each trial before being picked up, dried with absorbent wipes, and placed back into their warmed home cage.

On the first day of testing, mice were given a cue test with the platform positioned just above the surface of the water and a metal blue flag placed upon it for easy visual location of the platform. This test allows for detection of individual visual and swimming-related motor deficits and allows the mice to habituate to the task (climbing on the platform to escape the water). The platform was placed in a different location for each of the four trials with spatial cues removed by encirclement of the pool with a white plastic curtain.

On the next day, acquisition trials began with the hidden platform remaining in the same location (“NW quadrant”) for all trials/days and the curtain drawn back for visibility of the spatial cues. Several measures (distances to platform) and criteria to reach the platform (approximately 90% success rate, approximately 20 second latency to find platform) during the acquisition phases were recorded and achieved before mice were deemed to have learned the task. The performances of the four trials were averaged for each animal per day until criteria were met.

#### Open field test

Naive mice were individually introduced into one of eight adjacent open field arenas for 30 min and allowed to explore. Open field arenas consisted of custom made clear acrylic boxes (43 × 43 × 32h cm) with opaque white acrylic siding surrounding each box 45 × 45 × 21.5h cm to prevent distractions from activities in adjacent boxes. Activity was monitored with two CCTV cameras (Panasonic WV-BP334) feeding into a computer equipped with Ethovision XT 11.5 for data acquisition and analyses. A white noise generator (2325-0144, San Diego Instruments) was set at 65 dB to mask external noises and provide a constant noise level. Fluorescent linear strip lights placed on each of the four walls of the behavioral room adjacent to the ceiling provided a lower lighting (200 lux) environment than ceiling lighting to encourage exploration.

#### Contextual fear conditioning

A dedicated fear conditioning room in the TSRI Florida Mouse Behavior Core contains four fear conditioning devices that can be used in parallel. Each apparatus was an acrylic chamber measuring approximately 30 x 30 cm (modified Phenotyper chambers, Noldus, Leesburg, VA). The top of the chamber is covered with a unit that includes a camera and infrared lighting arrays (Noldus, Ethovision XT 11.5, Leesburg, VA) for monitoring of the mice. The bottom of the chamber is a grid floor that receives an electric shock from a shock scrambler that is calibrated to 0.40 mA prior to experiments. The front of the chamber has a sliding door that allows for easy access to the mouse. The chamber is enclosed in a sound-attenuating cubicle (Med Associates) equipped with a small fan for ventilation. Black circular, rectangular and white/black diagonal patterned cues were placed outside each chamber on the inside walls of the cubicles for contextual enhancement. A strip light attached to the ceilings of the cubicles provided illumination. A white noise generator (~65 dB) was turned on and faced toward the corner of the room between the cubicles. The fear conditioning paradigm consisted of two phases, training, followed by testing 1 and 26, or 30 d thereafter. The 4.5 min training phase consisted of 2.5 min of uninterrupted exploration. Two shocks (0.40 mA, 2 s) were delivered, one at 2 min 28 s, the other at 3 min and 28 s from the beginning of the trial. During testing, mice were placed into their designated chambers and allowed to roam freely for 5 min. Immobility durations (s) and activity (distances moved (cm)) during training and testing were obtained automatically from videos generated by Ethovision software. Activity suppression ratio levels were calculated: 0-2 min activity during testing/0-2 min activity during training + testing.

---

**Figure 2 - Supplement.**
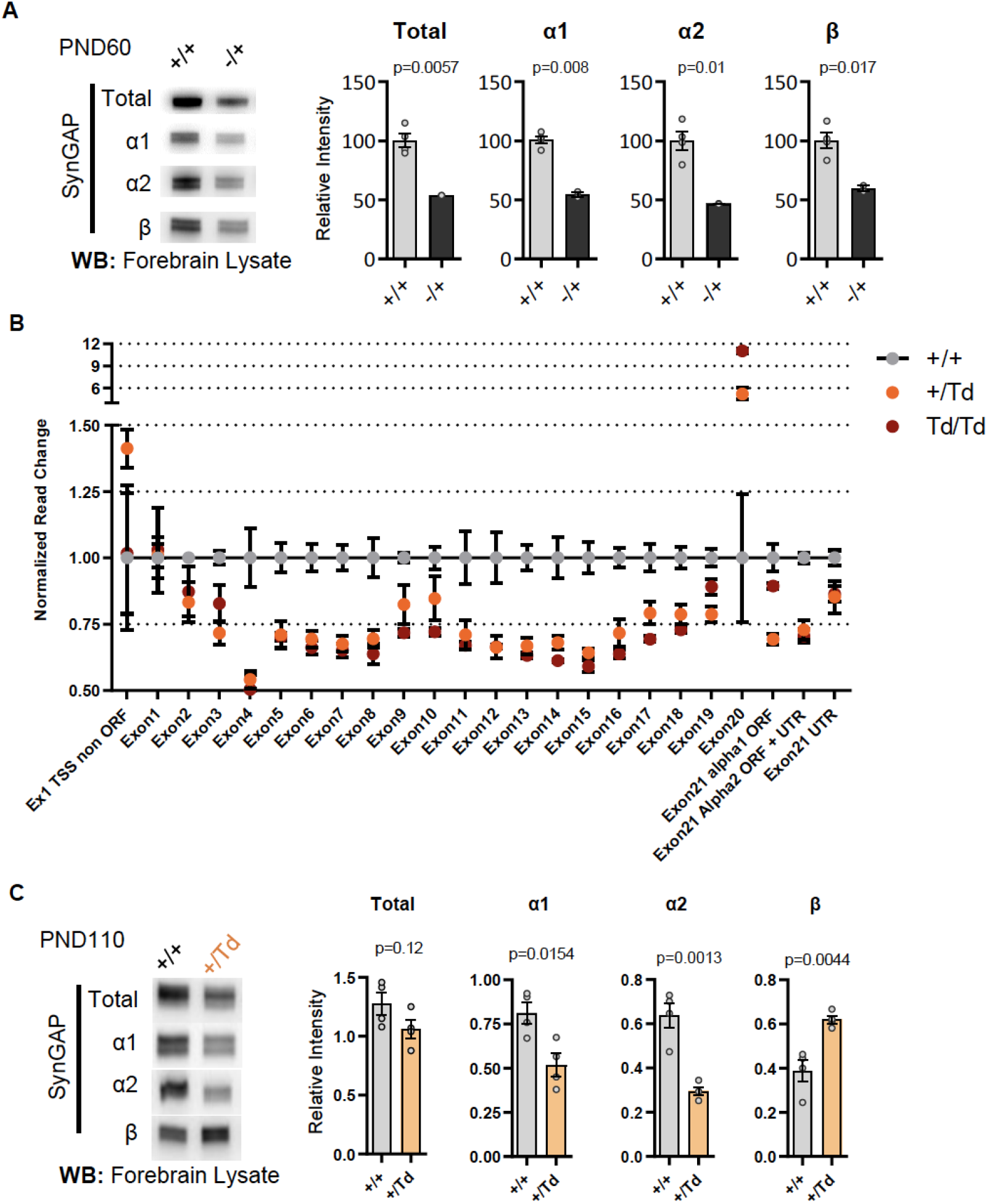
**(A)** Representative western blots demonstrating total SynGAP and isoform expression level in forebrain lysates from *Syngap1^+/+^* and *Syngap1^+/-^* mice. Relative intensity of bands normalized to total protein signal. Statistical significance is determined by unpaired t-test. Total: t(4)=5.403, α1: t(4)=9.044, α2: t(4)=4.473, β: t(4)=3.931 **(B)** *Syngap1* exon usage in +/+, +/Td, and Td/Td mice. **(C)** Representative western blots showing expression levels of total SynGAP and individual isoforms at PND110 from in +/+ and +/Td mice. Unpaired t-test. Total: t(6)=1.784, α1: t(6)=3.351, α2: t(6)=5.678, β: t(6)=4.425

**Figure 3 - Supplement.**
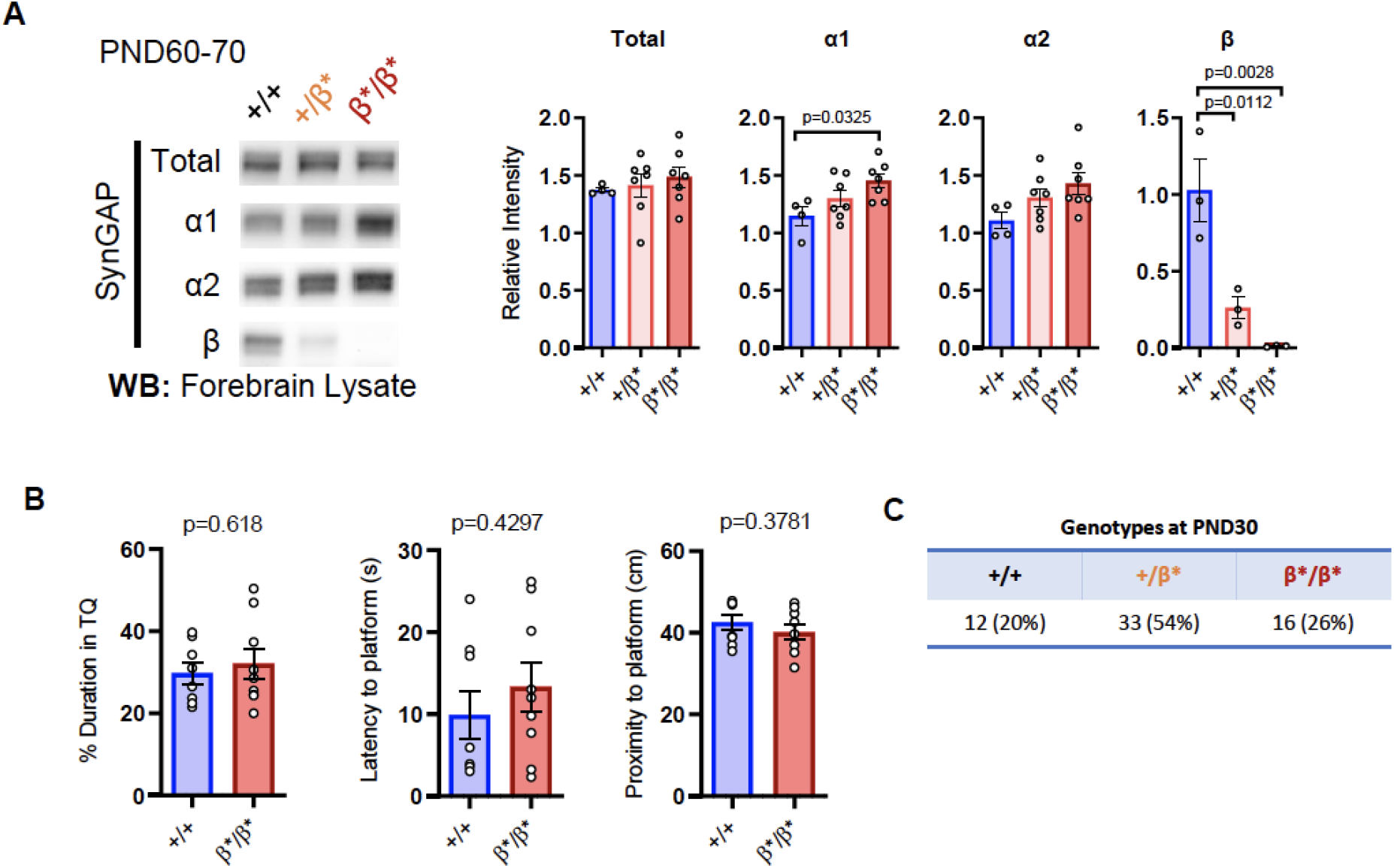
**(A)** Representative western blots showing expression levels of total SynGAP and individual isoforms at PND60-70. ANOVA with Tukey’s multiple comparisons test. Total: F(2, 15) = 0.3477, p=0.7119. α1: F(2, 15) = 4.102, p=0.0379. α2: F(2, 15) = 2.664, p=0.1023. β: F(2, 6) = 18.22, p=0.0028. **(B)** 24hr probe test in Morris water maze. Unpaired t-test. % Duration in target quadrant: t(15)=0.5093. Latency to platform: t(15)=0.8115. Proximity to platform: t(15)=0.9083 **(C)** Genotype numbers and ratios derived from heterozygous breeding of β* line (7 litters).

**Figure 4 - Supplement.**
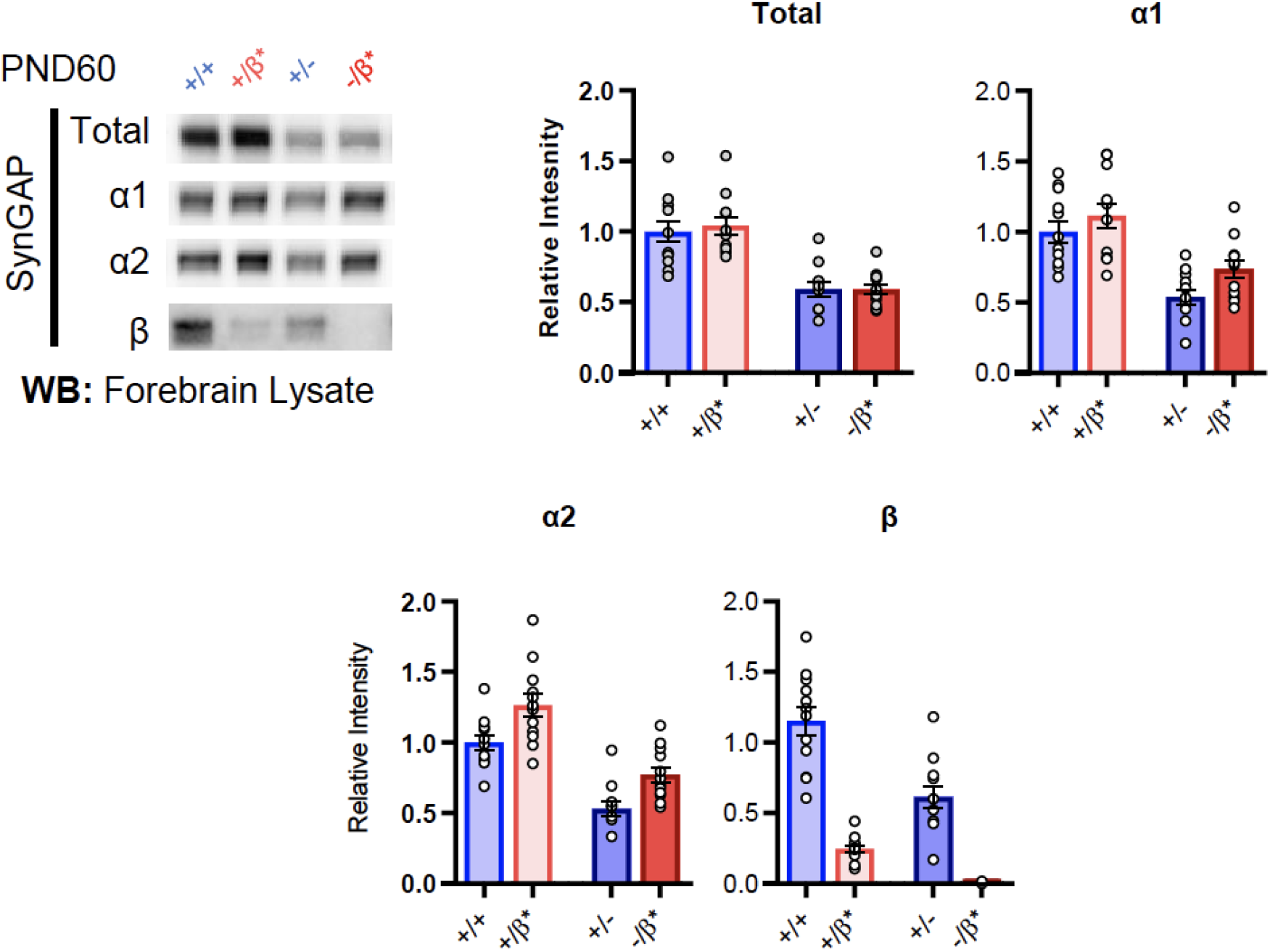
Representative western blots showing expression levels of total SynGAP and individual isoforms at PND60 for all genotypes. Two-way ANOVA with Tukey’s multiple comparison test. **Total:** (-) allele F(1, 44)=58.57, p<0.0001; β* allele F(1, 44)=0.1181, p=0.7327. Allelic Interaction F(1, 244)=0.1839, p=0.6701. **α1:** (-) allele F(1, 44)=35.37, p<0.0001; β* allele F(1, 44)=4.932, p=0.031; Allelic Interaction F(1, 44)=0.3615, p=0.5508. **α2:** (-) allele F(1, 44)=63.95, p<0.0001; β* allele F(1, 44)=18.00, p<0.0001; Allelic Interaction F(1, 44)=0.03486, p=0.8527. **β:** (-) allele F(1, 20)=9.149, p=0.0067; β* allele F(1, 20)=9.676, p=0.0055; Allelic Interaction F(1, 20)=0.3027, p=0.5883.

**Figure 5 - Supplementary.**
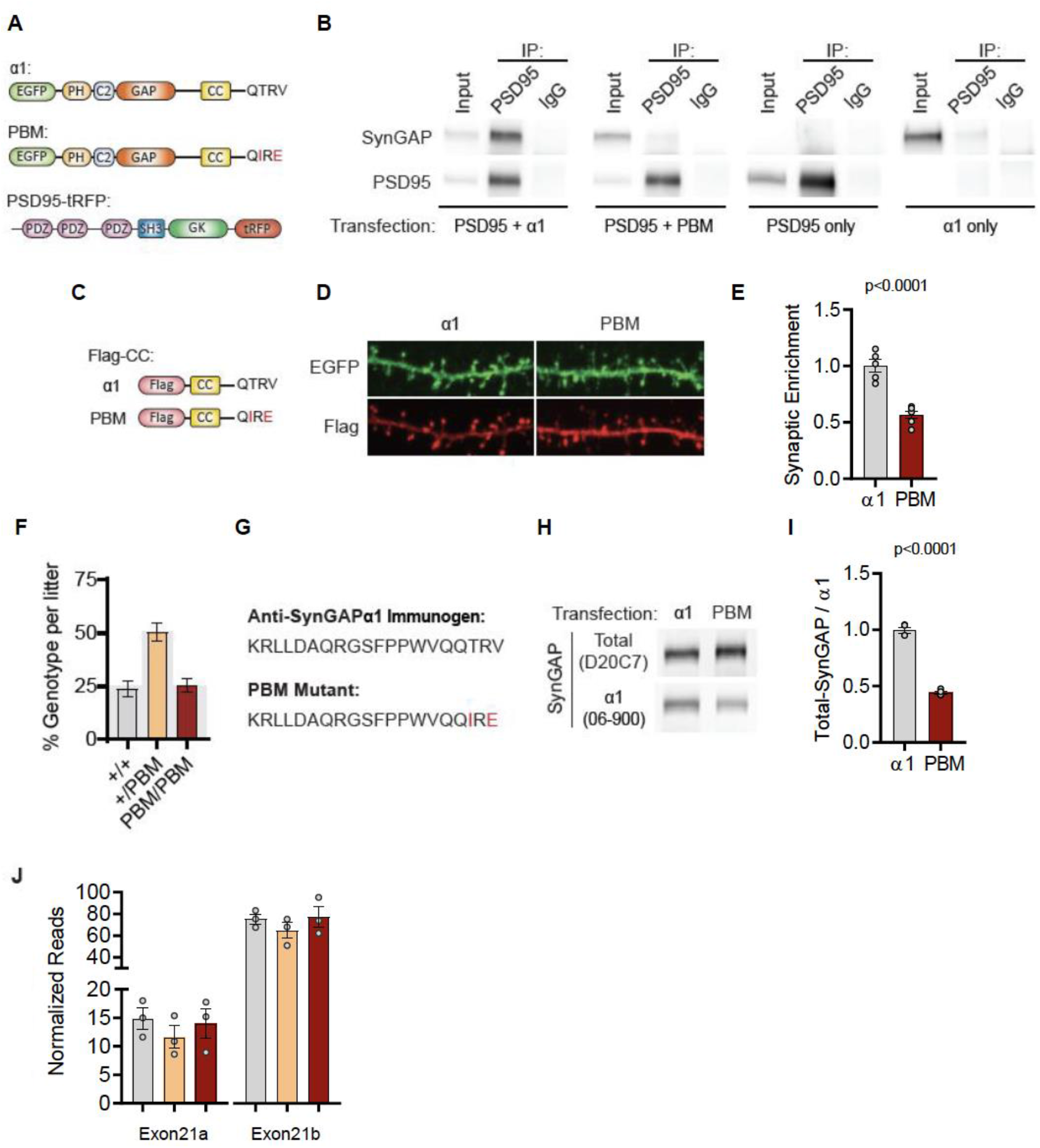
**(A)** Illustrations of constructs expressed in H293T cells to study PDZ-dependent interaction between SynGAP and PSD95. **(B)** Coimmunoprecipitation of PSD-95 and SynGAPα1 from transfected H293T cells. PSD95-tRFP coprecipitates with SynGAPα1. This Interaction was disrupted by PBM mutations. **(C)** Illustrations of Flag-tagged SynGAP C-terminal constructs expressed in primary cortical neurons. **(D)** Subcellular localization of wild-type or PBM mutated Flag-CCα1 in primary forebrain neurons. Note that Flag-CC α1 is heavily enriched in dendritic spines compared to Flag-CC PBM. Height of the image is 5μm. **(E)** Quantification of synaptic enrichment of Flag-CC constructs. Enrichment in dendritic spines were calculated as the ratio of Flag signal in spines vs dendrites over ratio of EGFP signal in spines vs dendrites. Unpaired t-test, t(9)=6.982 p<0.0001. Introduced point mutations impeded the enrichment of Flag-tagged SynGAPα1 C-terminal construct in primary forebrain neurons. **(F)** Genotype frequencies observed from 15 litters following heterozygous crosses. Expected mendelian ratio is highlighted with gray. **(G)** Antigen for α1-specific antibody in comparison to PBM mutant C-tail. **(H)** Reduced antigenicity of α1 antibody against PBM mutant C-terminus. H293T cells were transfected with either wild-type or PDZ-binding mutant form of EGFP-SynGAPα1. Lysates were probed for both Pan-SynGAP (D20C7) and α1-specific (06-800) antibody. Relative reduction in α1 to Pan-SynGAP signal demonstrates ~50% reduction in antigenicity. **(I)** Quantification of (D) Unpaired t-test. t(6)=19.16, n=4, p<0.0001. **(J)** SynGAP α1 mRNA levels in forebrain transcriptome. Normalized reads of Exon21b (specific to α1) were shown in linear scale. ANOVA F(2,6)=0.3009, n=3, p=0.7507. No significant changes were found across genotypes indicating that point mutations do not influence the mRNA expression levels.

## Notes

### Competing Interest Statement

The authors have declared no competing interest.

## References

1. Deciphering Developmental Disorders, S., Large-scale discovery of novel genetic causes of developmental disorders. Nature, 2015. 519(7542): p. 223–8.

2. Deciphering Developmental Disorders, S., Prevalence and architecture of de novo mutations in developmental disorders. Nature, 2017. 542(7642): p. 433–438.

3. Hamdan, F.F., et al., Mutations in SYNGAP1 in autosomal nonsyndromic mental retardation. N Engl J Med, 2009. 360(6): p. 599–605.

4. Vlaskamp, D.R.M., et al., SYNGAP1 encephalopathy: A distinctive generalized developmental and epileptic encephalopathy. Neurology, 2019. 92(2): p. e96–e107.

5. Parker, M.J., et al., De novo, heterozygous, loss-of-function mutations in SYNGAP1 cause a syndromic form of intellectual disability. Am J Med Genet A, 2015. 167a(10): p. 2231–7.

6. Mignot, C., et al., Genetic and neurodevelopmental spectrum of SYNGAP1-associated intellectual disability and epilepsy. J Med Genet, 2016. 53(8): p. 511–22.

7. Iossifov, I., et al., The contribution of de novo coding mutations to autism spectrum disorder. Nature, 2014. 515(7526): p. 216–21.

8. Satterstrom, F.K., et al., Large-Scale Exome Sequencing Study Implicates Both Developmental and Functional Changes in the Neurobiology of Autism. Cell, 2020. 180(3): p. 568–584 e23.

9. Holder, J.L., Jr., F.F. Hamdan, and J.L. Michaud, SYNGAP1-Related Intellectual Disability, in GeneReviews((R)), M.P. Adam, et al., Editors. 1993: Seattle (WA).

10. Weldon, M., et al., The first international conference on SYNGAP1-related brain disorders: a stakeholder meeting of families, researchers, clinicians, and regulators. J Neurodev Disord, 2018. 10(1): p. 6.

11. Llamosas, N., et al., SYNGAP1 Controls the Maturation of Dendrites, Synaptic Function, and Network Activity in Developing Human Neurons. J Neurosci, 2020. 40(41): p. 7980–7994.

12. Kilinc, M., et al., Species-conserved SYNGAP1 phenotypes associated with neurodevelopmental disorders. Molecular and Cellular Neuroscience, 2018. 91: p. 140–150.

13. Gamache, T.R., Y. Araki, and R.L. Huganir, Twenty Years of SynGAP Research: From Synapses to Cognition. J Neurosci, 2020. 40(8): p. 1596–1605.

14. Kim, J.H., et al., SynGAP: a synaptic RasGAP that associates with the PSD-95/SAP90 protein family. Neuron, 1998. 20(4): p. 683–91.

15. Chen, H.J., et al., A synaptic Ras-GTPase activating protein (p135 SynGAP) inhibited by CaM kinase II. Neuron, 1998. 20(5): p. 895–904.

16. Ozkan, E.D., et al., Reduced cognition in Syngap1 mutants is caused by isolated damage within developing forebrain excitatory neurons. Neuron, 2014. 82(6): p. 1317–33.

17. Araki, Y., et al., Rapid dispersion of SynGAP from synaptic spines triggers AMPA receptor insertion and spine enlargement during LTP. Neuron, 2015. 85(1): p. 173–89.

18. Clement, J.P., et al., Pathogenic SYNGAP1 mutations impair cognitive development by disrupting maturation of dendritic spine synapses. Cell, 2012. 151 (4): p. 709–23.

19. Clement, J.P., et al., SYNGAP1 links the maturation rate of excitatory synapses to the duration of critical-period synaptic plasticity. J Neurosci, 2013. 33(25): p. 10447–52.

20. Aceti, M., et al., Syngap1 Haploinsufficiency Damages a Postnatal Critical period of Pyramidal Cell Structural Maturation Linked to Cortical Circuit Assembly. Biological Psychiatry, 2015.

21. Michaelson, S.D., et al., SYNGAP1 heterozygosity disrupts sensory processing by reducing touch-related activity within somatosensory cortex circuits. Nature Neuroscience, 2018. 21(12): p. 1–13.

22. Araki, Y., et al., SynGAP isoforms differentially regulate synaptic plasticity and dendritic development. Elife, 2020. 9.

23. Gou, G., et al., SynGAP splice variants display heterogeneous spatio-temporal expression and subcellular distribution in the developing mammalian brain. J Neurochem, 2020. 154(6): p. 618–634.

24. McMahon, A.C., et al., SynGAP isoforms exert opposing effects on synaptic strength. Nat Commun, 2012. 3: p. 900.

25. Rumbaugh, G., et al., SynGAP regulates synaptic strength and mitogen-activated protein kinases in cultured neurons. Proceedings of the National Academy of Sciences of the United States of America, 2006. 103(12): p. 4344–4351.

26. Vazquez, L.E., et al., SynGAP regulates spine formation. J Neurosci, 2004. 24(40): p. 8862–72.

27. Komiyama, N.H., et al., SynGAP regulates ERK/MAPK signaling, synaptic plasticity, and learning in the complex with postsynaptic density 95 and NMDA receptor. J Neurosci, 2002. 22(22): p. 9721–32.

28. Sullivan, B.J., et al., Low-Dose Perampanel Rescues Cortical Gamma Dysregulation Associated With Parvalbumin Interneuron GluA2 Upregulation in Epileptic Syngap1(+/-) Mice. Biol Psychiatry, 2020. 87(9): p. 829–842.

29. Spicer, T.P., et al., Improved Scalability of Neuron-Based Phenotypic Screening Assays for Therapeutic Discovery in Neuropsychiatric Disorders. Mol Neuropsychiatry, 2018. 3(3): p. 141–150.

30. Kim, J.H., et al., The role of synaptic GTPase-activating protein in neuronal development and synaptic plasticity. J Neurosci, 2003. 23(4): p. 1119–24.

31. Yokoi, S., et al., 3’UTR Length-Dependent Control of SynGAP Isoform alpha2 mRNA by FUS and ELAV-like Proteins Promotes Dendritic Spine Maturation and Cognitive Function. Cell Rep, 2017. 20(13): p. 3071–3084.

32. Creson, T.K., et al., Re-expression of SynGAP protein in adulthood improves translatable measures of brain function and behavior. eLife, 2019. 8: p. e46752.

33. Guo, X., et al., Reduced expression of the NMDA receptor-interacting protein SynGAP causes behavioral abnormalities that model symptoms of Schizophrenia. Neuropsychopharmacology, 2009. 34(7): p. 1659–72.

34. Zeng, M., et al., Phase Transition in Postsynaptic Densities Underlies Formation of Synaptic Complexes and Synaptic Plasticity. Cell, 2016. 166(5): p. 1163–1175.e12.

35. Gou, G., et al., SynGAP Splice Variants Display Heterogeneous Spatio-Temporal Expression And Subcellular Distribution In The Developing Mammalian Brain. bioRxiv, 2019: p. 681148.

36. Li, J., et al., Spatiotemporal profile of postsynaptic interactomes integrates components of complex brain disorders. Nature Neuroscience, 2017. 20: p. 1150.

37. Wilkinson, B., J. Li, and M.P. Coba, Synaptic GAP and GEF Complexes Cluster Proteins Essential for GTP Signaling. Scientific Reports, 2017. 7(1): p. 5272.

38. Volianskis, A., et al., Different NMDA receptor subtypes mediate induction of long-term potentiation and two forms of short-term potentiation at CA1 synapses in rat hippocampus in vitro. The Journal of Physiology, 2013. 591 (4): p. 955–972.

39. Wang, C.-C., R.G. Held, and B.J. Hall, SynGAP Regulates Protein Synthesis and Homeostatic Synaptic Plasticity in Developing Cortical Networks. PLoS ONE, 2013. 8(12): p. e83941.

40. Hamdan, F.F., et al., De novo SYNGAP1 mutations in nonsyndromic intellectual disability and autism. Biol Psychiatry, 2011. 69(9): p. 898–901.

41. Berryer, M.H., et al., Mutations in SYNGAP1 cause intellectual disability, autism, and a specific form of epilepsy by inducing haploinsufficiency. Hum Mutat, 2013. 34(2): p. 385–94.

42. Lim, K.H., et al., Antisense oligonucleotide modulation of non-productive alternative splicing upregulates gene expression. Nat Commun, 2020. 11(1): p. 3501.

43. Zhang, L.I. and M.M. Poo, Electrical activity and development of neural circuits. Nature Neuroscience, 2001. 4: p. 1207–1214.

44. Lynch, G., C.S. Rex, and C.M. Gall, LTP consolidation: substrates, explanatory power, and functional significance. Neuropharmacology, 2007. 52(1): p. 12–23.

45. Dravid, S.M., et al., Subunit-specific mechanisms and proton sensitivity of NMDA receptor channel block. J Physiol, 2007. 581(Pt 1): p. 107–28.

